# TGF-β modulates cell fate in human ES cell-derived foregut endoderm by inhibiting multiple endogenous signaling pathways

**DOI:** 10.1101/2021.07.15.452587

**Authors:** Nina Sofi Funa, Kristian Honnens de Lichtenberg, Maria Skjøtt Hansen, Jonas van Cuyl Kuylenstierna, Kim Bak Jensen, Yi Miao, K. Christopher Garcia, Palle Serup

## Abstract

Genetic differences between pluripotent stem cell lines causes variable activity of extra-cellular signaling pathways, which limits the reproducibility of directed differentiation protocols. Here we used human embryonic stem cells (hESCs) to interrogate how exogenously provided factors modulate endogenous signaling events during specification of foregut endoderm lineages. We find that TGF-β1 activates an *OTX2*/*LHX1* gene regulatory network that promotes anterior fate by antagonizing endogenous Wnt signaling. In contrast to Porcupine inhibition, the effects of TGF-β1 cannot be reversed by exogenous Wnt ligands, suggesting that induction of SHISA proteins and intracellular accumulation of Fzd receptors make TGF-β1 treated cells refractory to Wnt signaling. Subsequently, TGF-β1-mediated inhibition of Bmp- and Wnt-signaling suppresses liver- and promotes pancreas fate. However, pancreas differentiation is delayed by TGF-β1-induced CYP26A1 expression and inhibition of RA signaling. Our study thus identifies multiple mechanisms of crosstalk between major developmental signaling pathways during foregut patterning.

## Introduction

Directed differentiation of human embryonic stem cells (hESC) seeks to recapitulate the signaling events that govern cell lineage decisions, from germ layer specification during gastrulation, to subsequent regionalization along the anterior-posterior (A-P) and dorso-ventral (D-V) axes. In vertebrates, a combination of Wnt and high Nodal signaling triggers formation of the definitive endoderm (DE) layer that emerges from the primitive streak. Closure of the DE along the anterior- and caudal intestinal portals leads to formation of the primitive gut (PG). The PG is patterned along the A-P and D-V axes to form three broad regions: foregut, midgut and hindgut that are further patterned into distinct organ territories, including thyroid, lung, liver, pancreas and the different subdivisions of the gastro-intestinal tract (Arnold and Robertson, 2009; Grapin-Botton, 2005; Tam and Loebel, 2007). As is also the case for meso- and ectoderm, posterior endoderm fates are promoted by high levels of Wnt, BMP and retinoic acid (RA) signaling (Bayha et al., 2009; Deimling and Drysdale, 2009; Spence et al., 2011; Stevens et al., 2017), whereas ligand-sequestering antagonists such as Dkk1, Sfrp, Noggin and Chordin, secreted from anterior visceral endoderm and anterior mesendoderm (ME), protect anterior tissues from Wnt and BMP signaling and thus permit the development of anterior fates (McLin et al., 2007; Rankin et al., 2011). Similarly, the retinoic acid (RA) catabolizing enzyme, Cyp26a1, expressed in the anterior epiblast and the anterior half of the newly formed germ layers, protects anterior structures from RA exposure (Abu-Abed et al., 2001; Abu-Abed et al., 2003; Ribes et al., 2007). Later, these signaling pathways are repurposed and act in a stage-dependent manner to promote distinct organ lineages including liver and pancreas (Rankin et al., 2018).

Liver and pancreas arise from common multipotent progenitors in the ventral foregut under the influence of FGF, BMP and Wnt signals emanating from adjacent, mesodermal tissues (Zaret, 2008). Dose-dependent FGF signaling from cardiogenic mesoderm and BMP-signaling from the septum transversum mesenchyme induces hepatic fate and suppresses pancreatic fate (Deutsch et al., 2001; Jung et al., 1999; Rossi et al., 2001; Serls et al., 2005). Accordingly, most hESC pancreas differentiation protocols use BMP signaling inhibitors to reduce liver specification (Nostro et al., 2011; Pagliuca et al., 2014; Rezania et al., 2014). Conversely, TGF-β signaling has been shown to favor pancreatic over hepatic specification in hESC cultures (Loh et al., 2014). However, it is not clear how TGF-β and BMP signaling interact during these fate decisions.

How Wnt signaling acts on the multipotent foregut progenitors to specify organ fate is not fully understood (reviewed in Zaret, 2008). Ectopic Wnt signaling in *Xenopus* foregut progenitors inhibits the development of foregut organ buds, including pancreas, liver and lung, while repression of Wnt signaling cause an expansion of liver and pancreas buds (McLin et al., 2007). However, this can be interpreted as Wnt repression being required for specification of anterior endoderm, and thus foregut progenitors. In zebrafish, loss of canonical Wnt2/Wnt2bb signaling prevents liver and swim bladder specification and the pancreas and anterior intestines expand (Ober et al., 2006; Poulain and Ober, 2011). More recent studies in mouse and human systems indicate that non-canonical Wnt signaling is able to promote pancreatic over liver fate choice, while contradictory results were found regarding the ability of canonical Wnt signaling to promote liver fate (Mahaddalkar et al., 2020; Rodriguez-Seguel et al., 2013). Thus, the interaction between signals that regulate segregation of liver and pancreas primordia are still not fully understood.

Similarly, we only have a limited understanding of how exogenously supplied growth factors affect endogenous signaling pathways at the various steps of hESC differentiation protocols. Although differences in endogenous signaling activity associated with different hESC lines (Ortmann et al., 2020) is an obvious source of variation in differentiation outcomes, the role of endogenous signaling pathways has not received a lot of attention. Generally, endogenously active pathways are inhibited by addition of antagonists, if considered detrimental to the desired differentiation outcome, often determined by trial-and-error approaches. Conversely, pathways whose activity is considered desirable are stimulated by addition of agonists. Yet, studies of embryonic development have uncovered numerous examples of crosstalk among signaling pathways and it is to be expected that variability of endogenous signaling, as well as addition of exogenous factors, will affect such crosstalk and thus the outcome of in vitro PSC differentiation in a cell line dependent manner.

Here we investigated how exogenously provided growth factors and inhibitors affect endogenous signaling events during early pancreatic lineage specification in differentiating hESC cultures. By screening major developmental signaling pathways we identified TGF-β1 and the porcupine inhibitor IWP-L6 as potent inducers of pancreatic fate when present during development of foregut endoderm and early pancreatic progenitors. We confirm the anteriorizing effect of Wnt inhibition and further show that TGF-β1 anteriorizes endoderm by inhibiting the expression and/or function of components in the Wnt signaling pathway. Concurrently, TGF-β1 and IWP-L6 both stimulate proliferation of OTX2^+^ anterior endoderm cells. We next show that Wnt/β-catenin signaling promotes liver fate and suppress pancreas fate in hESC cultures and that TGF-β1 suppresses liver fate by dual inhibition of BMP and Wnt signaling in foregut progenitors. Unexpectedly, we find that TGF-β1-induced CYP26A1 inhibits RA signaling to delay pancreas specification and maintain foregut progenitors in an OTX2^+^ state.

Our work uncovers mechanisms that underlie an extensive crosstalk between developmental signaling pathways in foregut progenitors as they develop towards liver and pancreas during hESC differentiation. Knowledge about such signaling dynamics and their effect on progenitor proliferation will enable more efficient in vitro generation of insulin-producing beta cells.

## Results

### TGF-β1 and Wnt inhibitors promote expression of pancreatic progenitor markers

When subjecting a *PDX1*^EGFP/+^ HUES4 reporter cell line (Ameri et al., 2017) to a control pancreas differentiation protocol (Rezania et al., 2014) we noted a disappointingly low expression of *PDX1* at the end of stage 5 (S5), where endocrine differentiation commences (Figure 1A and 1B). As induction of definitive endoderm (DE) appeared efficient at S1 (Figure S1A), we reasoned that subsequent anterior-posterior (A-P) patterning and/or specification of pancreatic endoderm (PE) was suboptimal in our cultures. We therefore added agonists or antagonists of selected pathways active in AP-patterning and pancreas specification to differentiating *PDX1*^EGFP/+^ HUES4 cells from S2 to S5 and monitored GFP expression at the end of S5 (D13). FACS analysis showed a significant increase in GFP^+^ positive cells when blocking Wnt-secretion with the porcupine inhibitor IWP-L6 or adding either recombinant TGF-β1 or Activin A, while perturbation of Notch signaling and relevant receptor tyrosine kinase pathways had only minimal effects (Figure 1B). Consistently, a prominent increase in PDX1 protein expression was seen by immunofluorescent (IF) staining and western blotting following treatment with IWP-L6 or TGF-β1 (Figure S1B and S1D). Gene expression analysis by qRT-PCR at D3-D13 showed that *PDX1* and *NKX6-1* transcripts increased from S3 and S5, respectively, after IWP-L6 treatment, while the effect of TGF-β1 on *PDX1* expression was only evident at S4. We also noted that expression of the anterior marker *OTX2* was significantly higher in both IWP-L6- and TGF-β1-treated samples than in controls at S2 and S3. Conversely, expression of the posterior marker *CDX2* and the early liver lineage marker alpha fetoprotein (*AFP)* was suppressed from S2 and onwards by both treatments (Figure 1C). These results suggest that IWP-L6 inhibits the secretion of posteriorizing Wnt proteins in our cultures, consistent with how Wnt proteins act *in vivo* (Loh et al., 2014; McLin et al., 2007). This notion is supported by IWP-L6 being effective when added selectively at S2/3, while later addition had no effect on pancreatic marker expression at S5 (Figure S1C).

**Figure 1.**
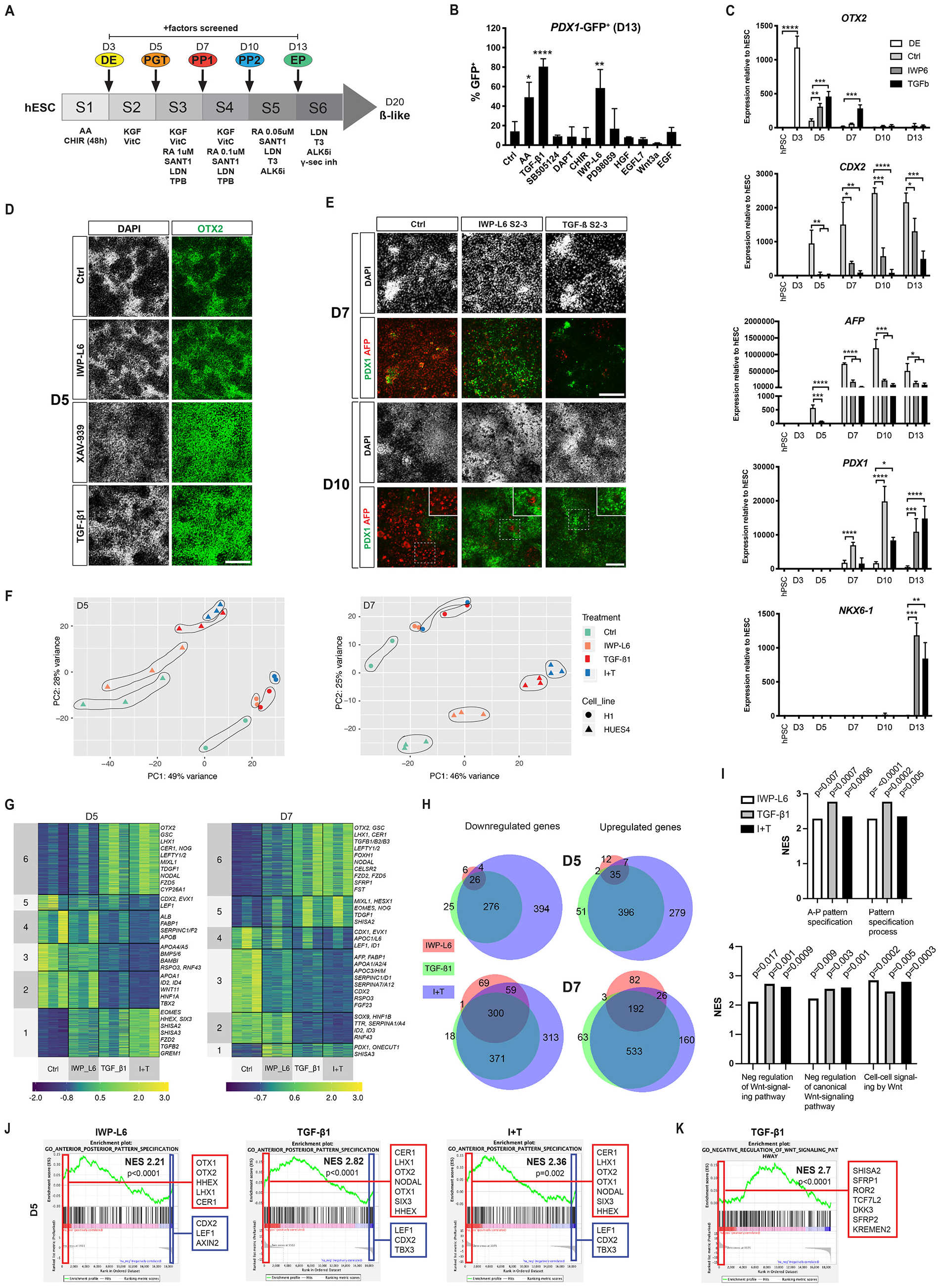
TGF-β1 and IWP-L6 anteriorize endoderm and promote pancreas over liver fate. (A) Schematic overview of the basic differentiation protocol. The stages when compounds being screened were added are indicated. DE: definitive endoderm; PGT: primitive gut tube; PP1: pancreatic progenitor 1; PP2: pancreatic progenitor 2; EP: endocrine precursor. (B) FACS analyses of *PDX1*^EGFP/+^ HUES4 cells showing percent GFP^+^ cells at D13 after treatment with indicated factors during S2-S5. Mean ± SD, N=3. * p < 0.05, ** p < 0.005, **** p < 0.0001. (C) qRT-PCR analyses of *OTX2*, *CDX2*, *AFP*, *PDX1* and *NKX6-1* expression in differentiating HUES4 cells at the indicated time points after treatment with the indicated factors during S2-S3. Data are shown relative to undifferentiated HUES4 cells. Mean ± SD, N=3, * p < 0.05, ** p < 0.005, *** p < 0.0005, **** p < 0.0001. (D) IF staining for OTX2 in differentiating HUES4 cells at D5 after treatment with the indicated factors during S2. The cells were counterstained with DAPI. Scale bar, 50 μm. (E) IF staining for PDX1 (green) and AFP (red) in differentiating HUES4 cells at D7 and D10 after treatment with the indicated factors during S2-S3. The cells were counterstained with DAPI. Insets show higher magnifications of boxed areas. Scale bars, 50 μm. (F) Principle component analysis (PCA) showing clustering along PC1 and PC2 for *PDX1*^EGFP/+^ HUES4 (n=3) and H1 (n=2) cells treated with IWP-L6, TGF-β1 or IWP-L6 + TGF-β1 (I+T) during S2 (D5) or S2-S3 (D7). (G) Heatmaps scaled by row to show patterns of regulated genes in *PDX1*^EGFP/+^ HUES4 cells (n=3) by k-means clustering analysis of RNA-seq data. Signature genes are indicated for each cluster. (H) Venn diagram showing overlap of all deregulated genes at D5 and D7 with an FDR <0.1 and fold change >1.5. (I) GSEA plots comparing gene expression data from each treatment at D5 (n=3) with the indicated gene sets. Normalized Enrichment Scores (NES) and p-values are indicated. (J-K) GSEA plots comparing D5 gene expression data from each treatment (J) or TGF-β1 (K) with gene sets for anterior-posterior pattern specification or negative regulation of Wnt signaling, respectively (N=3). Signature genes from leading and trailing edge analyses are shown in red and blue boxes, respectively. Normalized Enrichment Scores (NES) and p-values are shown. See also Figure S1.

Analysis of OTX2 protein expression confirmed a marked increase at S2 after treatment with IWP-L6 or the Tankyrase inhibitor XAV-939, supporting that inhibition of Wnt signaling anteriorizes the cells (Figure 1D and Figure S1D). Notably, TGF-β1 treatment increased the number of OTX2^+^ cells at S2 to the same extent as IWP-L6 (Figure 1D). Further IF analysis showed an increase in PDX1^+^ cells already at S3 following IWP-L6 treatment and at S4 following TGF-β1 treatment, both at the expense of AFP^+^ cells that were present in remarkably high numbers in control cells at both time points (Figure 1E and Figure S1D). Taken together, these results suggest that endogenous Wnt proteins posteriorize hESC-derived DE cultures while exogenous TGF-β1 has the opposite effect.

### TGF-β1 anteriorize endoderm by antagonizing Wnt signaling

To gain a better understanding of how IWP-L6 and TGF-β1 affect hESC-derived DE cultures, we performed RNA-seq analysis of cells treated with IWP-L6, TGF-β1 or both at the end of S2 (D5) and S3 (D7) in the human ES cell lines HUES4 and H1 (Table S1). Principle component analysis (PCA) showed that the samples clustered according to both cell line and treatment. Importantly, the cells demonstrated a clear transcriptional response to the different treatments and the trajectories of the clusters along PC1 and PC2 were similar in both cell lines (Figure 1F).

DESeq2 analysis found 92, 811 and 1417 genes differentially expressed (adjusted p-value < 0.05) in D5 cells treated with IWP-L6, TGF-β1, or both, respectively, compared to control cells. Similarly, 732, 1481, and 1804 genes were differentially expressed at D7. 56, 484, and 717 genes were upregulated (log2 FC > 1) and 36, 327, and 700 genes were downregulated (log2 FC < −1) at D5. At D7 303, 791, and 911 genes were upregulated and 429, 690, and 1043 genes were downregulated (Table S2). Analysis by k-means clustering revealed that the majority of the regulated genes overlapped between the conditions (Figure 1G, 1H and Table S2). Genes upregulated by TGF-β1 and TGF-β1 + IWP-L6 at D5 were found in clusters 1 and 6. Strikingly, IWP-L6 treatment also markedly upregulated cluster 1 genes, while cluster 6 genes were weakly stimulated or unchanged (Figure 1G and Table S2). Cluster 1 and 6 included members of the gene regulatory network (GRN) responsible for anterior specification downstream of Smad2/3 such as *EOMES*, *GSC*, *HHEX*, *LHX1* and *OTX2* as well as downstream targets *CER1, NOG, FZD5* and *SHISA2 (*from here on termed the *OTX2/LHX1* GRN). Similarly, genes downregulated by TGF-β1 and TGF-β1 + IWP-L6 at D5 (clusters 2 – 5) were typically also downregulated by IWP-L6, albeit more moderately. These included the posterior markers *EVX1* and *CDX2* as well as the Wnt and BMP agonists and target genes *WNT11*, *RSPO3*, *LEF1*, *RNF43* and *BMP5/6*, *BAMBI* and *ID2/4*, respectively (Figure 1G and Table S2). Gene set enrichment analysis (GSEA) was dominated by enrichment of Gene Ontology (GO) categories relating cell cycle (see below), but also terms related to A-P pattern specification and negative regulation of Wnt signaling were enriched (Figure 1I, Table S3 and Table S4). Leading edge analysis of the latter two gene sets identified genes in the *OTX2/LHX1* GRN, including *OTX2, HHEX, LHX1, SIX3*, *FZD5* and *CER1* (Figure 1J) and known Wnt pathway genes *SHISA2*, *SFRP1/2*, *ROR2*, *TCF7L2*, *KREMEN2* following TGF-β1 treatment (Figure 1K).

### TGF-β1 and IWP-L6 stimulate proliferation of anterior foregut progenitors

Addition of TGF-β1 and/or IWP-L6 during S2-3 resulted in a marked increase in cell density from D5 and onwards (Figure 2A and 2F). Consistently, proliferation markers such as *MKI67*, *PCNA*, *CCNA2*, *CCNB1*, *CDCA8, CDC20*, *CDC25A/B*, *E2F1*, and *MCM6* were upregulated at D5 by TGF-β1 and IWP-L6 treatment (Figure 2B and Table S2). The top ten GO terms at D5 all related to processes involved in cell division and GSEA of IWP-L6 and/or TGF- β1-treated D5 samples showed a significant enrichment of cell cycle related genes (Figure 2C, 2D, Table S3 and S4). We therefore assayed cell proliferation at D5 and D7 by EdU incorporation. An increase in proliferation was observed by FACS-analysis of EdU-incorporation at both timepoints (Figure 2E) and blocking TGF-β signaling with the ALK4/5 inhibitor SB505124 significantly reduced EdU-incorporation at D7 (Figure 2F). The increase in EdU-percentage following IWP-L6 was also reversed with SB505124, indicating that TGF-β1 and inhibition of Wnt-signaling might promote proliferation through a common mechanism. To test whether IWP-L6 and TGF-β1 promote proliferation of bona fide PFG cells and not simply favor the emergence of a new proliferating cell type, we quantified the percentage of OTX2^+^EdU^+^ cells at D5. IF analysis confirmed that both IWP-L6 and TGF-β1 showed a significant increase in the percentage of OTX2^+^EdU^+^ cells (Figure 2G and 2H).

**Figure 2.**
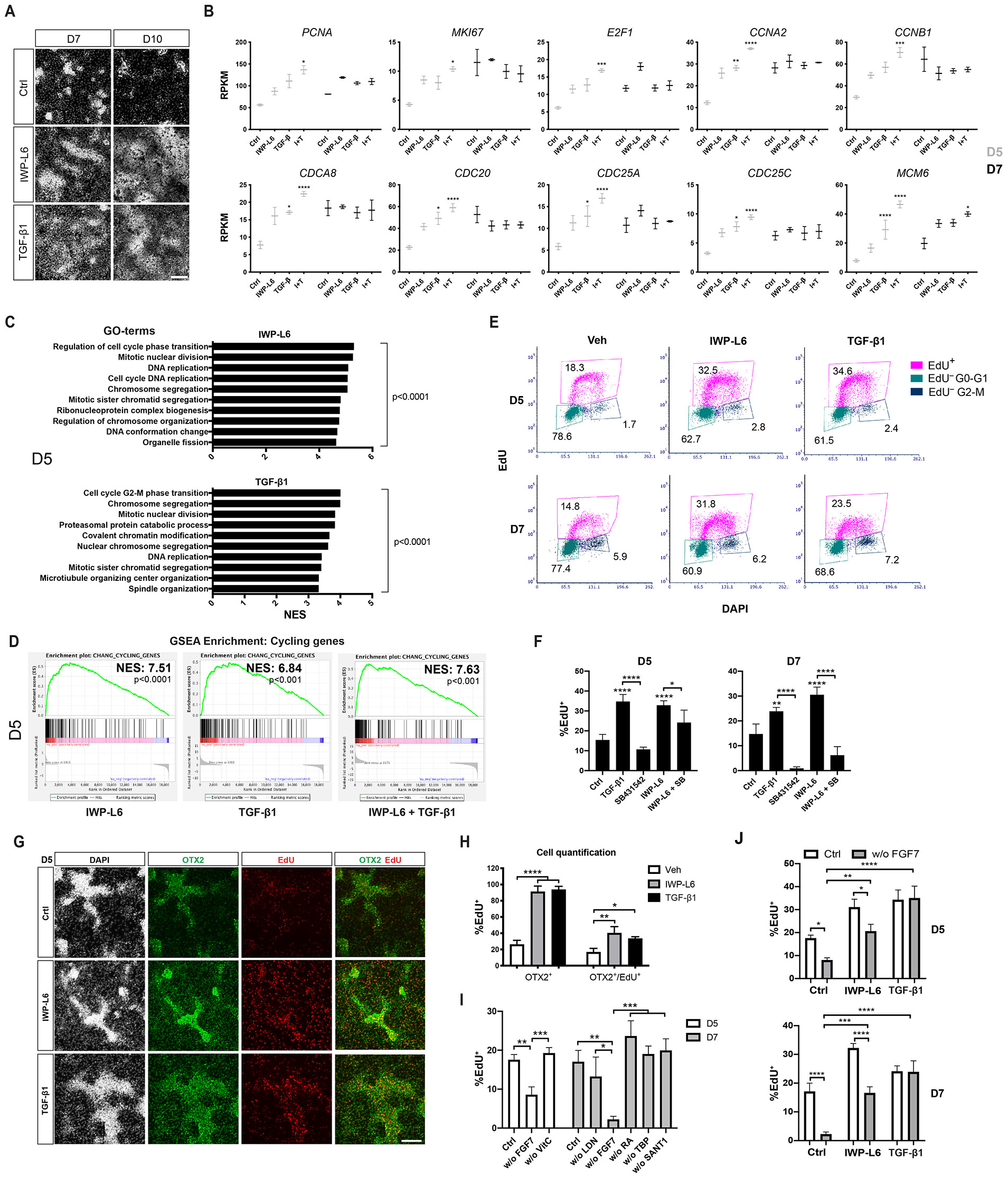
TGF-β1 and IWP-L6 stimulate proliferation of anterior foregut progenitors. (A) DAPI staining showing increased cellular density at D7 and D10 following treatment with IWP-L6 or TGF-β1 during S2-S3. Scalebar, 100μm. (B) Expression of cell cycle related genes by RNA-seq in *PDX1*^EGFP/+^ HUES4 cells treated with IWP-L6, TGF-β1 or IWP-L6 + TGF-β1 (I+T) during S2 (D5) or S2-S3 (D7). Mean ± SEM, N=3), * padj < 0.05, ** padj < 0.005, *** padj < 0.0005, **** padj < 0.0001. (C) NES-scores for the ten most enriched biological process GO terms for genes enriched at D5 in cells treated with IWP-L6 or TGF-β1 during S2. (D) GSEA plots comparing gene expression data from each treatment at D5 with a gene set for cycling genes. Normalized Enrichment Score (NES) and p-values are indicated. (E) FACS analysis of EdU incorporation and DNA content in differentiated *PDX1*^EGFP/+^ HUES4 cells at D5 and D7 after treatment with vehicle, IWP-L6 or TGF-β1 during S2 (D5) or S2-S3 (D7). (F) Percentage of EdU^+^ cells in differentiated *PDX1*^EGFP/+^ HUES4 cells at D5 and D7 after treatment with the indicated factors. Mean ± SD, N=3), * p < 0.05, ** p < 0.005, **** p < 0.0001. (G) IF analysis of OTX2^+^/EdU^+^ co-expression following treatment with vehicle, IWP-L6 or TGF-β1. Scalebar, 100μm. (H) Quantification of OTX2^+^/EdU^+^ co-expression following treatment with vehicle, IWP-L6 or TGF-β1. Mean ± SD, N=3), * p < 0.05, ** p < 0.005, **** p < 0.0001. (I) Percentage of EdU^+^ cells following depletion of individual factors from the control medium. Mean ± SD, N=3, * p < 0.05, ** p < 0.005, *** p < 0.0005. (J) Percentage of EdU^+^ cells following withdrawal of FGF7 in cells treated with IWPL6 or TGF-β1. Mean ± SD, N=3, * p < 0.05, ** p < 0.005, *** p < 0.0005, **** p < 0.0001. See also Figure S2.

We next examined whether other growth factors present in the media during S2-3 were providing proliferative cues under control conditions. We quantified EdU incorporation after depletion of single factors from the control S2 and S3 media. We found that depletion of FGF7, but not any other factor, from the media caused a two-fold decrease in the percentage of EdU^+^ cells at D5 and almost 10-fold reduction D7 (Figure 2I), consistent with the role of mesenchymal FGF10 as a proliferative signal for multipotent pancreatic progenitor cells in mice (Bhushan et al., 2001). We then tested whether FGF7 was required for the proliferative effects of IWP-L6 and TGF-β1. Strikingly, both IWP-L6 and TGF-β1 was capable of inducing proliferation in the absence of FGF7, albeit IWP-L6 was slightly less potent compared to TGF-β1 (Figure 2J).

As we found TGF-β1 induced the OTX2/LHX1 GRN and OTX2 is able to directly activate cell cycle genes (Bunt et al., 2012), we examined publicly available ATAC-seq and ChIP-seq data for chromatin accessibility, Smad2 binding, and binding of selected TFs from the *OTX2/LHX1* GRN to cell cycle genes in human ES cell cultures differentiating towards endoderm (Geusz et al., 2021; Tsankov et al., 2015). This analysis revealed that cell cycle genes upregulated by TGF-β1 all showed binding of OTX2 in their promoter regions at the DE stage (Figure S2), suggesting that OTX2 could be directly activating cell cycle genes in proliferating foregut progenitors. Taken together, these findings suggest that Wnt-inhibition and TGF-β1 stimulate proliferation of foregut progenitors via OTX2-mediated activation of the cell cycle.

### TGF-β1 activate an *OTX2/LHX1* GRN that antagonize Wnt signaling at multiple levels

To better understand how TGF-β1 antagonized Wnt signaling, we first analyzed expression of Wnt pathway agonists, antagonists and target genes in response to TGF-β1 treatment. We noted high expression of *WNT11* and moderate expression of *WNT3*, *WNT5A* and *WNT5B* in control cultures, while *RSPO3* was the only R-Spondin expressed (Figure S3A and Table S1). Notably, TGF-β1 treatment caused a significant reduction in expression of *WNT11* and *RSPO3* as well as the target genes *LEF1* and *RNF43*, while similar reductions after IWP-L6 treatment only reached significance for *LEF1* and *RSPO3* (Figure 3A). As expected, blocking TGF-β1 action with SB431542 rescued expression of *RSPO3* and *LEF1* at D5, but had no effect on IWP-L6-induced changes (Figure S3B). TGF-β1 treatment stimulated expression of both soluble and intracellular Wnt antagonists, including *SFRP1*, *SHISA2* and *SHISA3* at D5, while *SFRP5* was suppressed. Similarly, we found induction of *SFRP1*, *SFRP2*, *SHISA2* and *SHISA4* at D7 (Figure 3B and Table S2). *CER1* was also strongly induced at both timepoints, but may not antagonize Wnt signaling in humans (see discussion).

**Figure 3.**
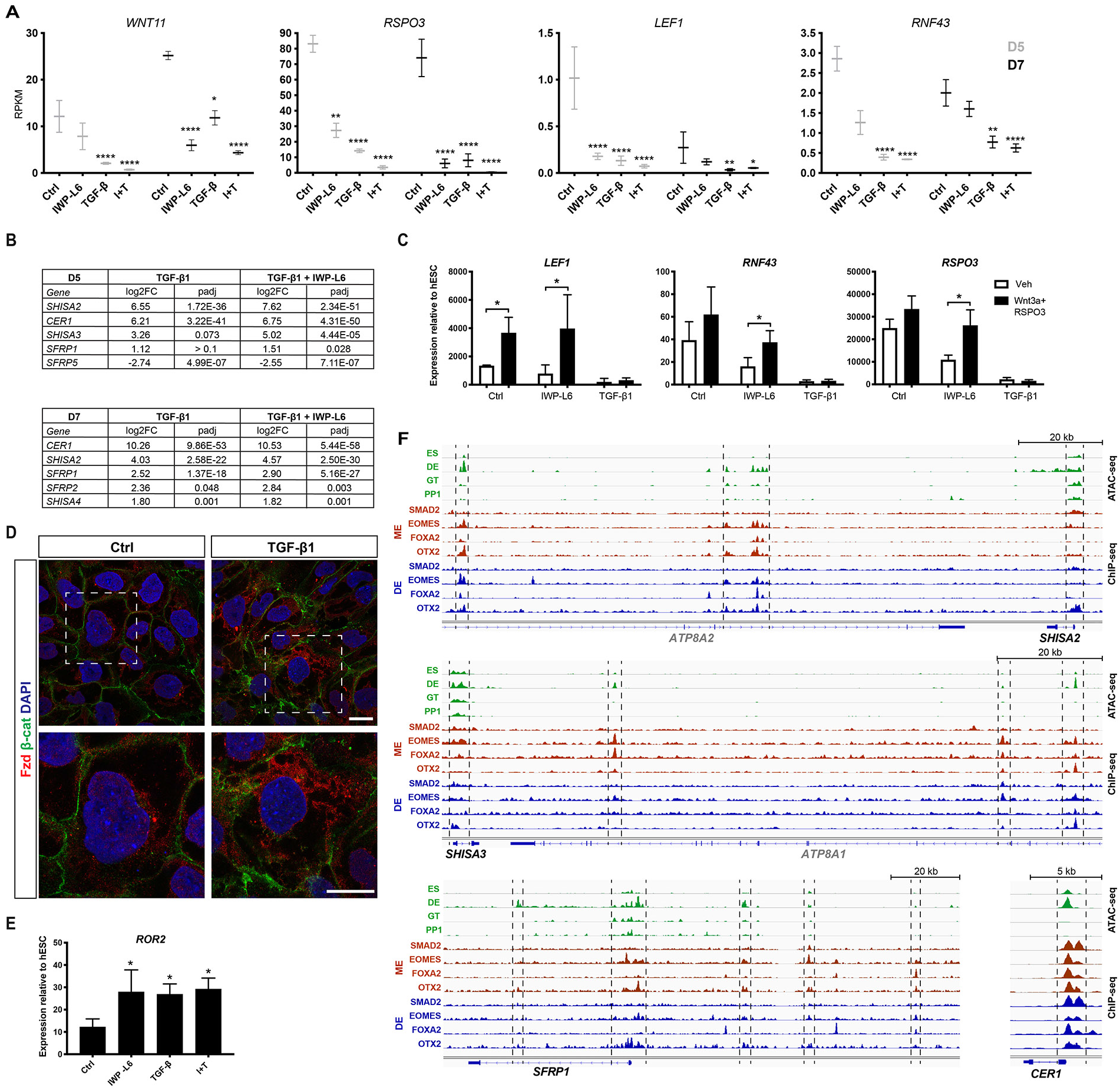
TGF-β1 activates an *OTX2*/*LHX1* GRN that antagonize Wnt signaling. (A) Expression of *WNT11, RSPO3, LEF1* and *RNF43* by RNA-seq in *PDX1*^EGFP/+^ HUES4 cells treated with vehicle (Ctrl), IWP-L6, TGF-β1 or IWP-L6 + TGF-β1 (I+T) during S2 (D5) or S2-S3 (D7). Mean ± SEM, N=3, * padj < 0.05, ** padj < 0.005, **** padj < 0.0001. (B) Differential expression of Wnt-signaling antagonists by DESeq2 analysis of RNA-seq data from *PDX1*^EGFP/+^ HUES4 cells treated with TGF-β1 or IWP-L6 + TGF-β1 during S2 (D5) or S2-S3 (D7). Log2 fold change (log2FC) relative to vehicle controls is shown as are adjusted p-values (padj). (C) qRT-PCR analysis of *LEF1, RNF43* and *RSPO3* expression at D5 in response to Wnt3a + RSPO3 stimulation during S2 in control cells (Ctrl) or cells treated with IWP-L6 or TGF-β1. Mean ± SD, N=3, * p < 0.05. (D) Confocal microscopy of D5 *PDX1*^EGFP/+^ HUES4 cells stained for β-catenin (green) and Fzd receptors (red) after treatment with vehicle (Ctrl) or TGF-β1 during S2. (E) qRT-PCR analysis of *ROR2* expression at D5 in *PDX1*^EGFP/+^ HUES4 cells treated with IWP-L6, TGF-β1 or IWP-L6 + TGF-β1 (I+T) during S2. Mean ± SD, N=3, * p < 0.05. (F) Signal tracks of ATAC-seq from Geusz et al. (2021) and SMAD2, EOMES, FOXA2 and OTX2 ChIP-seq data from Tsankov et al. (2015) at the *SHISA2*, *SHISA3*, *SFRP1* and *CER1* loci. Dashed lines are used to mark areas with differential chromatin accessibility at the DE stage, and TF binding at the ME and/or DE stages. ES: ES cells; DE: definitive endocerm; GT: primitive gut tube; PP1: pancreatic progenitor 1, ME: mesendoderm. See also Figure S3.

Since TGF-β1 treated D5 cells fail to activate Wnt target genes in response to exogenous Wnt3a/RSPO3 (Figure 3C), the induction of *SHISA2/3* caught our attention as the encoded proteins antagonize Wnt signaling cell autonomously by sequestering Frizzled (Fzd) receptors in the endoplasmatic reticulum (ER) (Onishi and Zou, 2017; Yamamoto et al., 2005). We therefore analyzed Fzd mRNA and protein expression in control, IWP-L6 and TGF-β1-treated cultures. The four highest expressed Fzd transcripts were *FZD4*, −*5,* −*7* and −*8* (RPKM > 10; Figure S3A and Table S1). Consistent with mouse *Fzd2*, *Fzd5*, *Fzd7* and *Fzd8* being members of the *Otx2/Lhx1* GRN (Costello et al., 2015; Sibbritt et al., 2018), treatment with IWP-L6 or TGF-β1 increased *FZD2* and *FZD5* expression significantly, with smaller, non-significant increases in *FZD7* and *FZD8* (Figure S3C and Table S2). To visualize FZD receptors and to address whether TGF-β1-induced SHISA2/3 expression correlated with FZD subcellular localization, we permeabilized control and TGF-β1-treated cells and labeled them with a human-Fc-tagged version of the next-generation surrogate Wnt; DRPB-Fz7/8, which recognizes Fzd1, −2, −5, −7, and −8 (Dang et al., 2019; Miao et al., 2020). Confocal imaging showed prominent intracellular accumulation of FZD-receptors in TGF-β1 treated samples compared to controls (Figure 3D), suggesting that SHISA2/3-mediated retention of FZD receptors in the ER may contribute to TGF-β1-induced suppression of Wnt signaling. When examining Wnt co-receptor expression we find a tendency to increased *ROR2* mRNA in the RNA-seq data in response to IWP-L6 and TGF-β1 (Figure S3D). We therefore analyzed *ROR2* expression by qRT-PCR and found a twofold increase (Figure 3E). Conversely, LRP4 was downregulated (Figure S3D), while the more highly expressed LRP5/6 remained unchanged (Table S2). Overall, these changes may bias any residual Wnt signaling towards the non-canonical pathway.

Studies in mice, fish and frogs have found that genes encoding signaling antagonists such as Cer1, Sfrp1 and Shisa2 as well as the Wnt receptors Fzd2, Fzd5, Fzd7 and Fzd8 are activated by the concerted action of Otx2 and Lhx1 (Costello et al., 2015; Fossat et al., 2015; Sibbritt et al., 2018), while ligands such as Wnt8 and Wnt11 are repressed by Gsc in concert with Otx2 (Seiliez et al., 2006; Yao and Kessler, 2001; Yasuoka et al., 2014). Since these TFs were strongly induced by TGF-β1 (Figure S3E), we assessed chromatin accessibility and TF binding for these Wnt pathway genes in human ES cell cultures differentiating towards endoderm (Geusz et al., 2021; Tsankov et al., 2015). This revealed enhanced chromatin accessibility and binding of *EOMES*, *FOXA2* and *OTX2* at two putative *cis*-acting regions located ~65 and ~130 kb downstream of the *SHISA2* gene at the DE stage (Figure 3F). Data for human LHX1 were not available, but Otx2 and Lhx1 have previously been shown to bind the mouse *Shisa2* gene (Costello et al., 2015). Similarly, two regions located ~30 and ~117 kb downstream of the *SHISA3* gene and multiple regions within an ~80 kb region upstream of *SFRP1* showed enhanced chromatin accessibility at the DE stage and binding of EOMES, FOXA2 and OTX2, while *CER1* showed enhanced chromatin accessibility and binding of SMAD2, EOMES, FOXA2 and OTX2 at the promoter region at both ME and DE stages (Figure 3F). Taken together, these findings suggest that TGF-β1 treatment at S2 acts by maintaining expression of a conserved *OTX2*/*LHX1*/*GSC* GRN that suppresses Wnt/β-catenin signaling via multiple mechanisms including ligand downregulation, induction of antagonists, and changes in co-receptor expression.

### TGF-β1 induced inhibition of BMP- and Wnt-signaling promotes pancreas over liver specification

The strong suppression of *AFP* expression by IWP-L6 and TGF-β1 prompted us to further investigate pancreas versus liver differentiation in our cultures. In addition to *AFP*, other early liver-specific genes such as*, APOA1, FABP1* and *TTR* were all reduced at D7 (Figure 4A). Furthermore, GSEA of the RNA-seq datasets showed that both IWP-L6 and TGF-β1 strongly reduced expression of liver specific genes (Figure 4B and S4A) and hepatocyte signatures were the most reduced among single cell RNA-seq-based cell type signatures during S2-3 (Figure S4B, Table S4 and S5). Furthermore, GSEA of our RNA-seq data compared to RNA-seq data from LASER-capture micro dissected dorsal pancreatic buds (DP) and hepatic cords (HC) of Carnegie stage 13 human embryos (Jennings et al., 2017) confirmed that expression of genes that were induced or reduced in IWP-L6-treated cells D7 corresponded to genes seen in the dorsal pancreas (DP high) and hepatic cord (HC high) gene sets, respectively (Figure 4C, S4C and Table S6). Notably, IWP-L6 or TGF-β1 treatment at S2 alone (D4-5) was sufficient to suppress AFP at D7, indicating that Wnt and TGF-β signaling modulate lineage segregation between liver and pancreas already at the primitive gut tube stage (Figure S4D). Taken together, these results indicate that canonical Wnt signaling promotes human liver specification, as it does in zebrafish (Ober et al., 2006; Poulain and Ober, 2011) and that TGF-β signaling has an opposing effect.

**Figure 4.**
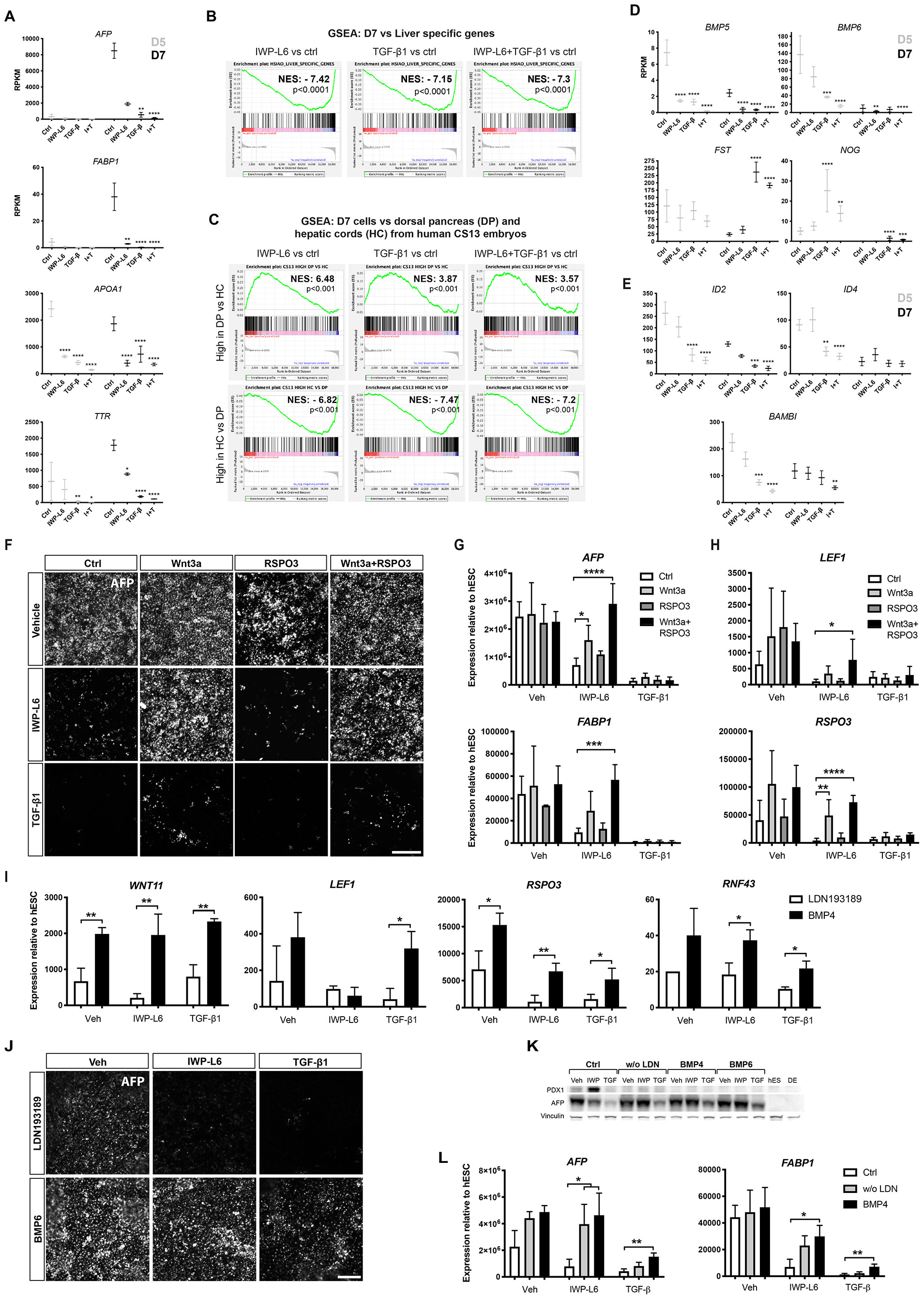
TGF-β1 promotes pancreas over liver fate by inhibiting BMP- and Wnt-signaling. (A) Expression of *AFP*, *APOA1*, *FABP1* and *TTR* by RNA-seq in *PDX1*^EGFP/+^ HUES4 cells treated with IWP-L6, TGF-β1 or IWP-L6 + TGF-β1 (I+T) during S2 (D5) or S2-3 (D7). Mean ± SEM, N=3, * padj < 0.05, ** padj < 0.005, **** padj < 0.0001. (B) GSEA plots comparing gene expression data from each treatment at D7 with a gene set for liver specific genes (Hsiao et al., 2001). Normalized Enrichment Score (NES) and p-values are shown. (C) GSEA plots comparing gene expression data from each treatment at D7 with genes highly expressed in human Carnegie Stage (CS)13 dorsal pancreas (DP) vs hepatic cords (HC) and genes highly expressed in HC vs DP (Jennings et al., 2017). (D-E) Expression of BMP ligands, secreted BMP antagonists (D) and BMP target genes (E) by RNA-seq in *PDX1*^EGFP/+^ HUES4 cells treated with IWP-L6, TGF-β1 or IWP-L6 + TGF-β1 (I+T) during S2 (D5) or S2-S3 (D7). Mean ± SEM, N=3, ** padj < 0.005, *** padj < 0.0005, **** padj < 0.0001. (F) IF analysis of AFP^+^ cells at D7 in response to Wnt3a, RSPO3 or Wnt3a + RSPO3 stimulation during S2-S3 in cells treated with vehicle, IWP-L6 or TGF-β1. (G-H) qRT-PCR analysis of *AFP*, *FABP1* (G), *LEF1* and *RSPO3* (H) expression at D7 in response to Wnt3a, RSPO3 or Wnt3a + RSPO3 stimulation during S2-S3 in *PDX1*^EGFP/+^ HUES4 cells treated with vehicle (Veh), IWP-L6 or TGF-β1. Mean ± SD, N=3, * p < 0.05. (I) Expression of *WNT11, LEF1*, *RSPO3* and *RNF43* by qRT-PCR in response to replacement of LDN193189 with BMP4 during S3 in *PDX1*^EGFP/+^ HUES4 cells treated with vehicle (Veh), IWP-L6 or TGF-β1. Mean ± SD, N=3, * p < 0.05, ** p < 0.005. (J) IF analysis of AFP^+^ cells at D7 in response to replacement of LDN193189 with BMP6 during S3 in *PDX1*^EGFP/+^ HUES4 cells treated with vehicle (Veh), IWP-L6 or TGF-β1. (K) Western blot showing AFP and PDX1 expression in *PDX1*^EGFP/+^ HUES4 cells cultured in standard S3 medium (Ctrl), S3 medium without LDN193189 (w/o LDN) or with replacement of LDN193189 with either BMP4 or BMP6 during S3, in cells treated with vehicle (Veh), IWP-L6 (IWP) or TGF-β1 (TGF). Also shown are hESC and DE stages. Vinculin is used as internal control. (L) Expression of *AFP* and *FABP1* by qRT-PCR in standard S3 medium (Ctrl), S3 medium without LDN193189 (w/o LDN) or with replacement of LDN193189 with BMP4 during S3, in cells treated with vehicle (Veh), IWP-L6 or TGF-β1. Mean ± SD, N=3, * p < 0.05, ** p < 0.005. See also Figure S4.

To understand how TGF-β signaling inhibits liver specification we interrogated our RNA-seq data for potential changes in BMP and Wnt signaling, two known liver promoting pathways. First, we found that the classical BMP ligands *BMP2*, *BMP4*, *BMP5*, *BMP6* and *BMP7* were expressed at moderate to high levels in control conditions (Figure S4E). TGF-β1 significantly suppressed expression of *BMP5* and *BMP6* and strongly induced the BMP antagonists *CER1*, *FST* and *NOG* (Figure 4D and S2E). In contrast, IWP-L6 had no effect on these genes with the exception of *BMP5*. Consistently, the downstream target genes *ID2*, *ID4* and *BAMBI* were downregulated by TGF-β1 but not by IWP-L6 (Figure 4E). Together, these findings suggest that TGF-β1-induced suppression of liver markers may be mediated, at least partly, through suppression of BMP signaling.

To test to whether reduced Wnt ligand availability might account for the IWP-L6- and TGF-β1-mediated suppression of liver markers, we co-treated cells with recombinant Wnt3a and/or RSPO3 during S2-3 and assayed *AFP* and *FABP1* expression. As expected, IWP-L6-mediated inhibition of liver markers D7 was fully reversed by co-treatment with Wnt3a with or without additional RSPO3 but not by RSPO3 alone (Figure 4F and 4G). However, Wnt3a, with or without RSPO3, did not prevent TGF-β1-induced inhibition of *AFP* and *FABP1* expression, indicating that suppression of Wnt ligand expression cannot fully account for the effect of TGF-β1 on liver differentiation seen at D7. As also seen at D5, treatment with Wnt3a and RSPO3 failed to induce expression of Wnt target genes in the presence of TGF-β1 (Figure 4H), showing that TGF-β1 blocks Wnt-signaling downstream of ligand availability. This may be due to increased expression of the antagonists *SFRP1/2* and *SHISA2/4* at D7 (Figure 3B).

Lastly, we asked whether BMP signaling influenced TGF-β1-mediated suppression of Wnt signaling and liver differentiation. Notably, BMP-signaling increased expression of Wnt11 and Wnt downstream targets *LEF1*, *RSPO3* and *RNF43*, and both IWP-L6- and TGF-β1-mediated suppression of these genes at D7 was partially rescued by replacing LDN193189 with BMP4 (Figure 4I). Strikingly, omitting the BMP receptor inhibitor LDN193189 and/or adding exogenous BMP4 or BMP6 counteracted IWP-L6- and TGF-β1-induced suppression of *AFP* and *FABP1* expression at D7, albeit the effect was most pronounced for IWP-L6 (Figure 4J–L). These data suggest that TGF-β1-mediated suppression of BMP signaling may contribute to reduced Wnt signaling by attenuating Wnt ligand expression.

### TGF-β1 delays pancreatic differentiation by attenuating RA signaling

To begin to unravel the mechanism responsible for the delayed induction of PDX1 following TGF-β1 treatment we interrogated our RNA-seq datasets for changes in signaling pathways known to be important for *PDX1* gene expression. We noted a strong induction of *CYP26A1* expression at D5 in TGF-β1 treated cells, which was augmented by IWP-L6 co-treatment, while IWP-L6 alone had no effect (Figure 5A). At D7 *CYP26A1* expression was high regardless of TGF-β1 addition, most likely due to the presence of 1 μM RA in S3 medium (Rezania et al., 2014). Nevertheless, we did observe reduced expression of *ALDH1A1* (encoding the RA-synthesizing enzyme RALDH1) at D7 in response to TGF-β1 treatment (Figure 5A). These data suggest that TGF-β1-treatment during S2 might desensitize the cells to RA added at S3. To test this hypothesis, we examined expression of pancreatic markers in control, TGF-β1 and IWP-L6-treated cells at D7, after co-treatment with the CYP26A1-inhibitor, R115866 at S2 (Figure 5B). Notably, addition of R115866 during S2 prevented the TGF-β1-induced suppression of *PDX1*, *SOX9*, *ONECUT1* and *HNF1B* expression and blunted the induction of *OTX2* expression at D7 (Figure 5C). However, inhibition of CYP26A1 did not prevent TGF-β1-mediated suppression of *AFP* expression. The effect of R115866 on OTX2 and PDX1 expression was also evident on the protein level (Figure 5D and 5E). To begin to understand how anterior *CYP26A1* expression is activated we again examined publicly available ATAC-seq and ChIP-seq data for chromatin accessibility and binding of Smad2 and TFs from the *Otx2/Lhx1* GRN to the *CYP26A1* locus. This analysis revealed regions of increased chromatin accessibility at D3 where cells have been exposed to Activin A and again at D7 where cells have been exposed to RA (Figure 5F). Notably, the immediate 5’-flanking region was bound by SMAD2 and EOMES at the ME stage and by OTX2 at the DE stage (Figure 5F). Taken together, these results suggest that TGF-β1 signaling delays pancreatic differentiation via SMAD2-induced *CYP26A1* expression, which desensitizes foregut cells to RA exposure.

**Figure 5.**
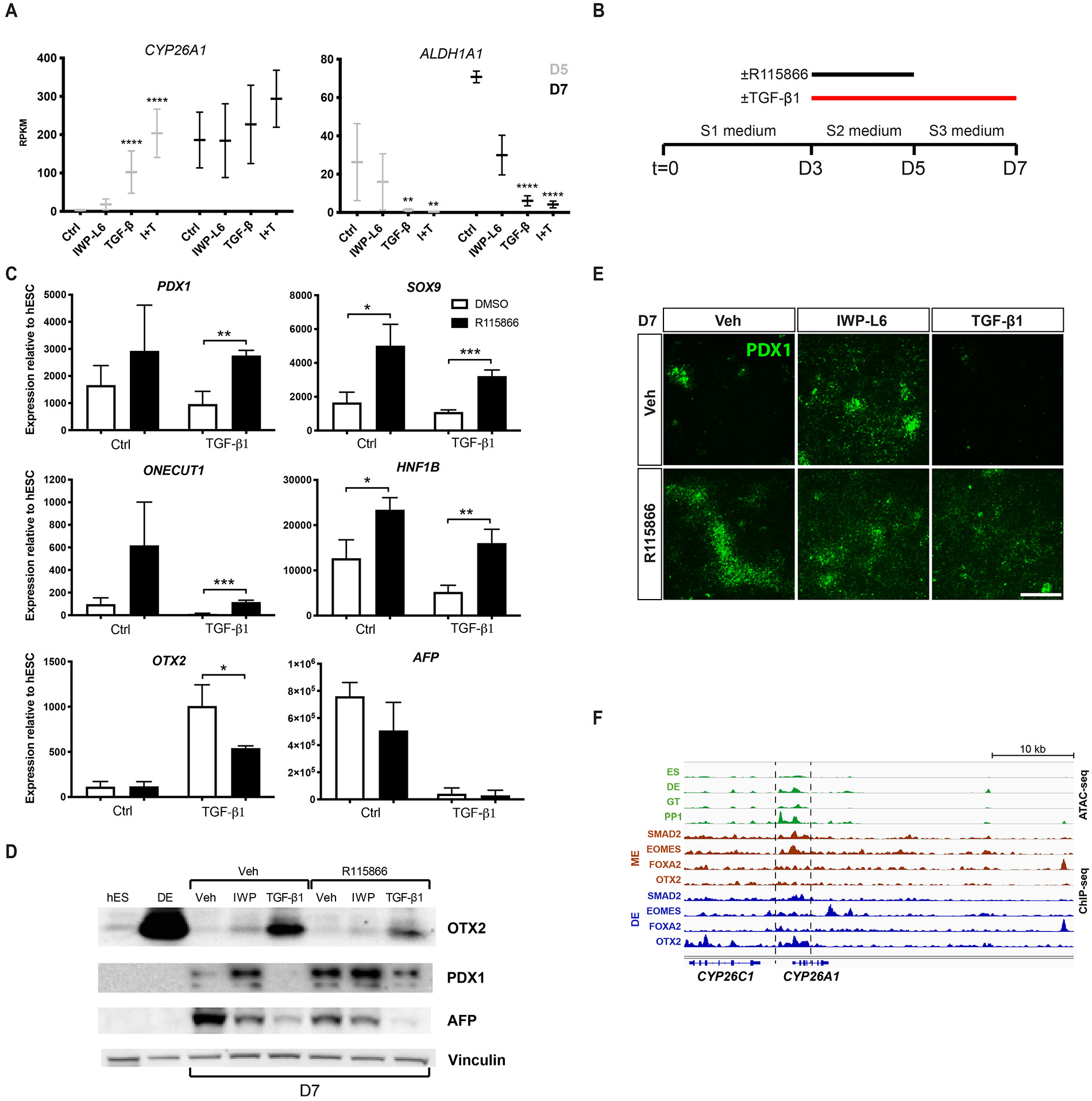
TGF-β1 delays pancreatic differentiation by attenuating RA signaling. (A) Expression of *CYP26A1* and *ALDH1A1* measured by RNA-seq in *PDX1*^EGFP/+^ HUES4 cells treated with IWP-L6, TGF-β1 or IWP-L6 + TGF-β1 (I+T) during S2 (D5) or S2-S3 (D7). Mean ± SEM, N=3, ** padj < 0.005 and **** padj < 0.0001. (B) Schematic overview of protocol using a CYP26-selective inhibitor, R115866 with TGF-β1. (C) Expression of *PDX1*, *SOX9*, *ONECUT1*, *HNF1B*, *OTX2* and *AFP* measured by qRT-PCR in D7 *PDX1*^EGFP/+^ HUES4 cells treated with vehicle (DMSO), TGF-β1 and/or R115866 as indicated. (D) Western blot showing OTX2, PDX1 and AFP expression in D7 *PDX1*^EGFP/+^ HUES4 cells treated with vehicle (Veh), IWP-L6, TGF-β1 and/or R115866 as indicated. Also shown are hES and DE stages. Vinculin is used as internal control. (E) IF analysis of PDX1 expression in D7 *PDX1*^EGFP/+^ HUES4 cells treated with vehicle (Veh), IWP-L6, TGF-β1 and/or R115866 as indicated. (F) Signal tracks of ATAC-seq from Geusz et al. (2021) and SMAD2, EOMES, FOXA2 and OTX2 ChIP-seq data from Tsankov et al. (2015) at the *CYP26A1* locus. Dashed lines are used to mark areas with differential chromatin accessibility at the DE and PP1 stages, and TF binding at the ME and/or DE stages. ES: ES cells; DE: definitive endoderm; GT: primitive gut tube; PP1: pancreatic progenitor 1, ME: mesendoderm.

## Discussion

Here we show that TGF-β signaling anteriorize hESC-derived definitive endoderm and subsequently promote pancreas fate over liver ditto via inhibition of Wnt/β-catenin and Bmp signaling in foregut progenitors. Our results uncover mechanisms used by TGF-β to modulate the activity of other signaling pathways operating in differentiating hESCs. We observed a very similar response to TGF-β1 and the Porcupine inhibitor IWP-L6, for which the most parsimonious explanation is suppression of Wnt signaling by TGF-β1. Remarkably, we also noted a strong stimulation of proliferation of S2 and S3 progenitors with these treatments. The exact mechanism still needs to be elucidated, but may involve direct activation of cell cycle genes by OTX2. Notably, the proliferative effect is independent of exogenous FGF. Together these observations should prove useful in relation to improving the efficacy of directed differentiation protocols for hESCs.

TGF-β/Nodal-induced Smad2/3 signaling promotes anterior fate in mice by inducing expression of Eomes that directly activates Foxa2 and the LIM domain homeobox transcription factor Lhx1, which in partnership with Otx2, activates numerous anterior ME genes as well as negative regulators of Bmp-, Nodal-, and Wnt/β-catenin signaling (Costello et al., 2015; Fossat et al., 2015; Ip et al., 2014; Sibbritt et al., 2018). We observed strong, induction of *EOMES*, *GSC*, *LHX1* and *OTX2* and other anterior TFs after TGF-β1 treatment at S2, indicating anteriorization of the endoderm formed at S1. Many of these markers were also induced by IWP-L6 treatment, as expected when inhibiting the posteriorizing effect of Wnt signaling. Together, these observations are best explained by suppression of Bmp-, Nodal and Wnt-signaling. This suppression appears to occur at several levels including repression of ligand expression and activation of antagonists such as *CER1*, *FST*, *LEFTY1/2, SFRP1* and *SHISA2/3*. The antagonists are well known targets of the Otx2/Lhx1 GRN in mouse and Xenopus (Costello et al., 2015; Fossat et al., 2015; Sibbritt et al., 2018) and our analysis of published ChIP-seq data (Tsankov et al., 2015) indicate that this may also be the case in humans. Future ChIP-seq analysis of LHX1 in human cells could bolster this notion. The mechanism underpinning the observed suppression of BMP and WNT ligand expression is unresolved, but GSC is a strong candidate as a direct repressor of ligand genes as it is induced by TGF-β1 in our cells and serve such a role in vivo, in concert with OTX2 (Yasuoka et al., 2014). Indeed, this study suggests that co-binding of OTX2 with either LHX1 or GSC likely determines activation or repression of OTX2 target genes, respectively, in *Xenopus* anterior development. SIX3, which in addition to anterior neural plate is expressed in hESC-derived ME and in early mouse DE (Shim et al., 2020), may also be involved. It is induced, albeit at low levels, by TGF-β1 in our cells and has been shown to repress BMP and WNT ligand expression in anterior neural plate (Gestri et al., 2005; Lagutin et al., 2003). Again, future ChIP-seq analyses of GSC and SIX3 in hESC-derived DE will help test these notions. The identity of the endogenous Wnt ligand(s) that act as posteriorizing factors in our cultures is not fully resolved by our data. However, we propose that Wnt11 is the key ligand for several reasons. First, Wnt11 is by far the highest expressed ligand in our culture with 6-10-fold higher RPKM values than any other Wnt ligand. Second, we find that Wnt11, together with RSPO3, is strongly suppressed by TGF-β1 treatment. Third, Wnt11 has been shown to be involved in foregut patterning in *Xenopus*, where Wnt11 activity must be suppressed by Sfrp5 in the anterior endoderm in order to maintain anterior foregut endoderm identity (Li et al., 2008; McLin et al., 2007). These studies found Sfrp5 expressed in surface cells of the foregut epithelium, in close proximity to the deep endoderm that expresses Wnt11, which activated both the Wnt/β-catenin and the Wnt/PCP pathways. Notably, morpholino-mediated depletion of Sfrp5 or Wnt11 overexpression in *Xenopus* embryos both caused a loss of foregut identity.

Looking at organ-specific markers, we found that liver markers were suppressed by both TGF-β1 and IWP-L6 and this was evident from D5 and onwards. Conversely, differentiation towards the pancreatic lineage was promoted, but only from D10 and onwards after TGF-β1 treatment. Analysis of endogenous signaling activity based on expression of well-known target genes at D5 and D7 revealed the expected suppression of Wnt signaling by IWP-L6 and suppression of both Wnt and Bmp signaling in response to TGF-β1. The TGF-β and BMP pathways often cross repress each other, through mechanisms that are not fully resolved, but may in some cases involve sequestration of a limited pool of Smad4 (Candia et al., 1997; Galvin et al., 2010). A reciprocal relationship between TGF-β and BMP in relation to induction of liver versus pancreas lineages has previously been reported in hESCs (Loh et al., 2014), but a mechanism was not described. Our results are thus consistent with previous observations and our RNA-seq data suggest that TGF-β1 may suppress Bmp signaling in foregut progenitors by repressing *BMP6* expression and inducing Bmp antagonists such as NOG, FST and CER1. *CER1* was strongly induced at both D5 and D7, while *NOG* and *FST* were induced at D5 and D7, respectively. While Bmp signaling is well known to promote liver development in vivo (Chung et al., 2008; Rossi et al., 2001), a mechanism involving suppression of BMP signaling may still appear surprising at first glance as the Bmp receptor inhibitor LDN193189 is present in S3 medium. However, as LDN193189 is only added at S3, it may leave time for endogenously produced BMPs to act at S2. Furthermore, LDN193189 only inhibits BMPR1B ~50% at the concentration used here (Sanvitale et al., 2013).

Notably, we found that blocking Wnt secretion via Porcupine inhibition promoted pancreas differentiation, while suppressing liver differentiation and that TGF-β1 also suppressed Wnt signaling at D7. In zebrafish, the ligands Wnt2 and Wnt2bb act via Fzd5 to activate Wnt/β-catenin signaling and promote liver development (Ober et al., 2006; Poulain and Ober, 2011), but whether and how Wnt/β-catenin signaling promotes mammalian liver development has been less clear. Non-canonical Wnt signaling has been reported to favor pancreas over liver fate, but no evidence of Wnt/β-catenin signaling promoting liver fate was found in the same study (Rodriguez-Seguel et al., 2013). However, recent work on hESC-derived foregut progenitors did find that Wnt/β-catenin signaling promoted liver fate (Mahaddalkar et al., 2020), and in agreement with this finding, we could rescue liver fate in IWP-L6 treated cultures by addition of Wnt3a. In our hESC cultures, the two most highly expressed *WNT* genes are *WNT5A* and *WNT11*, making these candidates for inducing liver fate, likely augmented by RSPO3. Like Wnt5a, Wnt11 is often considered a non-canonical ligand but both ligands can also activate the canonical pathway if the responding cells express the proper receptors (Li et al., 2008; Mikels and Nusse, 2006; Tao et al., 2005). However, the identity of a putative liver specifying Wnt ligand in vivo remains unknown, but candidates are Wnt2, Wnt2b and Wnt5a, which are all expressed in the mesoderm proximal to the developing liver (McMahon and McMahon, 1989; Monkley et al., 1996; Rodriguez-Seguel et al., 2013; Zakin et al., 1998). Mouse *Wnt2*/*Wnt2b* double mutants display normal liver and pancreas development (Goss et al., 2009), but additional Wnt ligands (e.g. Wnt5a) have been suggested to fulfill the liver specifying role in mammals (Poulain and Ober, 2011). Notably, the canonical co-receptor Lrp5 is enriched in liver progenitors compared to pancreas further suggesting that Wnt/β-catenin signaling may promote liver fate. A contributing factor to TGF-β1-mediated suppression of liver fate may be the upregulation of *ROR2* expression. In mouse foregut progenitors *Ror2* is expressed in pancreas-, but not liver progenitors and, as mentioned above, non-canonical signaling favors pancreas over liver fate (Rodriguez-Seguel et al., 2013).

As also seen at D5, TGF-β1 suppressed Wnt activity at D7 even in the presence of exogenous Wnt ligand, possibly by stimulating *SFRP1/2* and *SHISA2/4* expression at D7. *CER1* is also strongly induced, but human CER1, which is ~69% identical to mouse Cer1, may not antagonize Wnt signaling (Belo et al., 2000; Piccolo et al., 1999). Importantly, omitting LDN193189 from in the S3 medium, or replacing it with Bmp ligand, reactivated *WNT11*- and Wnt target gene expression and partially restored liver differentiation in the presence of TGF-β1. This suggests that Bmp signaling acts upstream of Wnt signaling and that TGF-β1-induced suppression of Wnt signaling could be mediated, at least partly, by inhibition of Bmp stimulated Wnt activity. Consistent with this notion, Bmp2b acts prior to Wnt2bb and Wnt2 in zebrafish liver development (Chung et al., 2008; Poulain and Ober, 2011). Taken together, our findings show that BMP- and Wnt/β-catenin signaling coordinate induction of liver lineage in human progenitors and suggest mechanisms for TGF-β1-mediated suppression of both pathways that ultimately promotes pancreatic over liver fate.

One notable difference between the effect of TGF-β1 and IWP-L6 was the delay in onset of pancreatic marker expression in TGF-β1 treated samples. Intriguingly, a previous study found suppression of Pdx1 expression in the developing pancreas after treatment of 3-4 and 5-6 somite stage half-embryo cultures with TGF-β2 and an increase after treatment of 5-6S half-embryo cultures with the ALK4,5,7 inhibitor SB431542. The underlying mechanism was not elucidated, but it was shown that changes in proliferation rate or apoptosis were not involved. (Wandzioch and Zaret, 2009). Importantly, our RNA-seq data showed that addition of TGF-β1 during S2 induced a profound increase in the expression of the RA-degrading enzyme CYP26A1. This induction is likely a reflection of the normal anterior expression pattern observed for *Cyp26a1* in mouse embryos (Abu-Abed et al., 2001; Abu-Abed et al., 2003; Ribes et al., 2007), and may be a direct induction by SMAD2, possibly in conjunction with EOMES. As RA is well-known inducer of Pdx1 and pancreatic fate (Kraus and Grapin-Botton, 2012; Micallef et al., 2005; Molotkov et al., 2005; Stafford and Prince, 2002), elevated CYP26A1 levels could explain the delay in the onset of pancreatic marker expression. Indeed, the delay was prevented by adding the CYP26A1 inhibitor R115866. This finding may partly explain why many pancreas differentiation protocols rely on ALK4,5,7 inhibitors during later stages (Nostro et al., 2011; Pagliuca et al., 2014; Rezania et al., 2014; Rezania et al., 2011).

In summary, our study suggests several mechanisms by which Bmp, TGF-β and Wnt signaling interact to control differentiation of hESC-derived foregut progenitors. Our findings improve the understanding of human foregut patterning and organ development and provide valuable new avenues for improving directed differentiation protocols for hESCs in order to obtain clinically relevant cell types in unlimited quantities.

## Acknowledgements

We thank Silvia Raineri for bioinformatics assistance and Jutta Bulkescher, Gelo de la Cruz, Paul van Dieken, Helen Neil, and Magali Michaut and the DanStem Research Platforms for technical assistance and the use of instruments. This work was supported by the Juvenile Diabetes Research Foundation International (3-APF-2017-390-A-N), the European Commission’s 7^th^ Framework Programme for Research (agreement 602587), the NIH (R01DK115728), Howards Hughes Medical Institute and Mathers Foundation. The Novo Nordisk Foundation Center for Stem Cell Biology is supported by grant number NNF17CC0027852.

## Author Contributions

N.S.F. and P.S. conceived the study, designed and interpreted experiments and wrote the manuscript. N.S.F, M.S.H. and J.v.C.K. carried out experiments, while N.S.F and K.H.L. performed the bioinformatics analyses. K.B.J., Y.M. and K.C.G. designed and produced the NGS detection reagent. All authors revised and approved the manuscript.

## STAR METHODS

### CONTACT FOR REAGENT AND RESOURCE SHARING

#### Lead contact

Further information and requests for resources and reagents may be directed to, and will be fulfilled by the lead contact, Palle Serup

#### Materials availability

This study did not generate new unique reagents.

#### Data and code availability

Sequencing datasets generated in this paper are available at ArrayExpress: E-MTAB-10715.

### EXPERIMENTAL MODEL AND SUBJECT DETAILS

#### Cell lines and culture conditions

The human PSC lines H1 (WA01, WiCell; RRID: CVCL_9771) and *PDX1*^EGFP/+^ HUES4 obtained from our facility (Ameri et al., 2017) were maintained in DEF-CS culture media (Takara Biosciences) following manufacturer’s instructions with daily media change and passaged every 3-4 days with TrypLE Express Enzyme (Thermo Fisher). All cells were cultured in a humidified 37°C, 5% CO_2_ incubator.

### METHOD DETAILS

#### Differentiation of hESCs in chemically defined conditions

The hESCs were differentiated to pancreatic progenitor cells by a previously described protocol (Rezania et al., 2014) with minor modifications. 150.000cells/cm2 were seeded in 6- or 24-well multi-well plates or in 8-well Ibidi chamber slides after single cell suspension for 3-5 min at 37 °C in TrypLE Express Enzyme (Thermo Fisher). After 48h, at day 0 (~90% confluency), the cells were washed once in 1× DPBS without Mg^2+^ and Ca^2+^ (Thermo Fisher) before addition of S1 basal media (MCDB131, Thermo Fisher) supplemented with Sodium Bicarbonate (1.5g/l, Sigma-Aldrich), Glucose (10mM, Sigma-Aldrich), GlutaMax (1x, Thermo Fisher), BSA (0,5%, Proliant Biologicals), CHIR-99021 (3μM, Axon Medchem) and Activin A (100 ng/ml, Peprotech). S1 cells (D3) were washed once in DPBS without Mg^2+^ and Ca^2+^ and further supplemented for 2d with S2 media including Sodium Bicarbonate (1.5g/l), Glucose (10mM), GlutaMax (1x), BSA (0,5%), L-Ascorbic Acid (0.25mM, Sigma-Aldrich) and recombinant human FGF7 (KGF) (50ng/ml, Peprotech). During S3-S4, the media was supplemented with Sodium Bicarbonate (2.5g/l), Glucose (10mM), GlutaMax (1x), BSA (2%), L-Ascorbic Acid (0.25mM), KGF (S3 50ng/ml and S4 2ng/ml), RA (S3 1μM and S4 0.1μM, Sigma-Aldrich), SANT-1 (0,25μM, Sigma-Aldrich), TPB (S3 200nM and S4 100nM, Sigma-Aldrich), LDN-193189 (S3 100nM and S4 200nM, Stemgent) and ITS-X (1:200, Thermo Scientific). During S5, the media was supplemented with Sodium Bicarbonate (1.5g/l), Glucose (20mM), GlutaMax (1x), BSA (2%), RA (0.05μM), SANT-1 (0.25mM), LDN-193189 (100nM), T3 (1μM, Sigma-Aldrich), ALK5i-II (10μM, Millipore), Zinc sulfate (10μM, Sigma-Aldrich), Heparin (10μM, Sigma-Aldrich) and ITS-X (1:200). In the initial screen, the media was further supplemented with the following factors during S2-4; Activin A (100ng/ml), TGF-β1 (10ng/ml, Peprotech), SB505124 (10 μM, Sigma-Aldrich), DAPT (10 μM, Selleckchem), CHIR-99021 (3μM), IWP-L6 (5μM, Axon Medchem), PD98059 (1 μM, Sellechchem), HGF (50 ng/ml, Peprotech), EGF-L7 (50ng/ml, Peprotech), Wnt3a (100 ng/ml, RnD Systems) or EGF (50ng/ml, Peprotech). In the following experiments, IWP-L6 (5μM) and/or TGF-β1 (10ng/ml) were added during S2-3 unless otherwise indicated. When applicable, SB431542 (10 ⍰M, Sigma-Aldrich), XAV939 (1⍰M, Sigma-Aldrich), R115866 (10 ⍰M, Sigma-Aldrich), BMP4/6 (50ng/ml, Peprotech), Wnt3a (100ng/ml) or RSPO3 (100ng/ml, Peprotech) were added during S2-3 according to experimental setup.

#### Flow cytometry

*PDX1*^EGFP/+^ HUES4 hESCs were differentiated S1-S4 and dissociated at D13 with TrypLE Express Enzyme and washed twice in DPBS without Mg^2+^ and Ca^2+^ + 2%BSA. Resuspended cells were further dissociated in tubes with cell strainer caps (Fisher Scientific), stained with DAPI and analyzed on an LSR Fortessa flow cytometer. FACS gating was determined using undifferentiated cells and non-GFP expressing cells.

#### Immunofluorescence analysis, Fzd detection and Imaging

Cells destined for immunofluorescence and confocal microscopy were grown on 24-well plates or 8-well Ibidi μ-slides (Ibidi), respectively. The cells were washed once in DPBS and fixed in 4% Formaldehyde (VWR) for 30 minutes, then permeabilized using 0.5% Triton-X in DPBS for 10 min at room temperature and blocked in SuperBlock (Thermo Fisher) for 30 minutes.

Primary antibodies (see Key Resources Table) were diluted in 0.1% Triton X-100 in DPBS. Incubation was done at 4°C overnight followed by 3×5 minutes wash in DPBS. Secondary antibodies (1:500, raised in Donkey) conjugated to either Alexa Fluor 488, Cy3 or Cy5 (all Jackson ImmunoResearch) were incubated for 45 minutes at room temperature and then washed 3×5min in DPBS followed by nuclear staining with DAPI (Thermo Fisher). FZD receptor expression was detected using a human Fc-tagged version of the NGS Wnt ligand: DRPB-Fz7/8 (Miao et al., 2020). Control and TGF-β1 treated cells, grown on 8-well Ibidi μ-slides (Ibidi), were fixed, permeabilized and incubated for 1h in media conditioned with cells expressing DRPB-Fz7/8 together with a primary antibody against β-catenin (BD Biosciences). Detection of DRPB-Fz7/8 was done with a PE-tagged anti-human Fc antibody (Jackson ImmunoResearch).

Images were captured and processed on a Zeiss Axioobserver using Plan-Apochromat 10x/0.45 and Plan-Apochromat 20x/0.8 objectives and ZEN software. Confocal images were captured on a Zeiss LSM780 confocal microscope using a Plan-Apochromat 63x/1.40 Oil objective. Figures were prepared using Adobe Photoshop CS6 and Adobe Illustrator CS6 (Adobe Systems, San Jose, CA, USA).

#### RNA extraction and quantitative real-time PCR

Total RNA was extracted with the RNeasy Plus Mini kit (Qiagen) and reverse-transcribed using the SuperScript III First-Strand synthesis kit (Invitrogen/Thermo Fisher). Quantitative real-time PCR experiments were performed using the StepOnePLus system (Applied Biosystems) and PowerUP SYBR Green Master Mix (Applied Biosystems/ThermoFisher). See Table S7 for primers. Relative changes in gene expression was compared to undifferentiated hESCs using the ΔΔCt method.

#### Western blotting

Harvested cells were lysed in RIPA buffer containing 1x phosphatase inhibitor cocktail (Sigma-Aldrich) and cOmplete Ultra Protease inhibitor (Thermo Scientific/Roche) on ice for 10min. Cell lysates were sonicated 5×30sec ON/OFF on a Diagenode BioRuptor in 1.5mL eppendorff tubes followed by centrifugation at 21 000g for 30 minutes at 4°C and saving the supernatant. Pierce BCA protein kit (ThermoFisher) was used to measure protein concentration on a Nanodrop 2000 (ThermoFisher). Lysates were boiled for 5 minutes in Laemmli sample buffer and 20-40 μg protein was separated by electrophoresis on NuPage 4-12% BisTris SDS-PAGE gels in MOPS buffer (Thermo Fisher) and transferred to PVDF membranes (Bio-Rad) using the BioRad Mini-Protean transfer system. Membranes were blocked in SuperBlock (Thermo Fisher) for 1 hour at room temperature and incubated with primary antibodies (see Key Resources Table) overnight at 4 °C. After three washes with TBS-T (0.1% Tween-20 in 1x Tris-buffered saline), the blot was incubated with respective secondary HRP antibodies at room temperature for 30 minutes. ECL Prime Western Blotting Detection Reagent was used for detection according to the manufacturer’s instruction (Sigma-Aldrich). For re-blotting, antibodies were stripped by Restore Western Blot Stripping Buffer (Thermo Fisher).

#### EdU-incorporation, analysis and quantification

*PDX1*^EGFP/+^ HUES4 cells were labeled with EdU and detected according to the Click-iT EdU Alexa Fluor 594 Flow Cytometry Assay kit (Thermo Fisher) protocol at D5 and D7 of differentiation. In brief, 10 1M EdU was added to the differentiation culture medium and cells were harvested and dissociated in to a single-cell suspension after 3h. The cells were fixed and permeabilized followed by EdU detection with the Click-iT EdU reagent and nuclear staining with DAPI. FACS gating was determined using non-EdU treated cells and EdU-treated cells without detection reagent. Cells were analyzed using a LSR Fortessa flow cytometer. Alternatively, EdU-treated cells were fixed, permeabilized and stained according to the Click-iT Plus EdU Cell Proliferation Kit for Imaging, 647 dye (ThermoFisher).

OTX2^+^ and OTX2^+^EdU^+^ cells were quantified from confocal images acquired on an LSM780 confocal microscope as described above. Cells were identified using the Spot function in Imaris™ (Bitplane) with the diameter set to 4.15 μm and the Co-localization function with the maximum distance between spots set to ≤2 μm to identify co-expressing cells.

#### RNA-seq

RNA-seq libraries were built using biological triplicates from *PDX1*^EGFP/+^ HUES4 cells and duplicates from H1 cells with 1μg of RNA using the NEB NEXT Ultra II RNA Library Prep Kit (NEB #E7770) and mRNA magnetic isolation module for poly(A) purification (NEB #E7490) with 5 cycles of amplification. Quality of the RNA and subsequently of the libraries were measured on a Fragment Analyser and the libraries were loaded accordingly and barcoded with NEB Next multiplex Oligos for Illumina (E7335 and E7500) on Illumina NextSeq 500 with Hi-output 1×75bp kit.

FASTQ-files were generated using bcl2Fastq (Illumina) and aligned to hg38 human genome using STAR (Dobin et al., 2013) with standard alignment settings resulting in >90% alignment of reads. Quantified gene count matrix with STAR (flag: --quantMode GeneCounts) and loaded into R using DESEQ2 package for differential expression analysis on gene level (Love et al., 2014). We performed quality control assessment including Principal Component Analysis after regularized logarithmic transformation. For the differential expression analysis for each cell line on each day we tested control (ctrl) vs. IWP-L6, TGF-β and IWP-L6+TGF-β (I+T). From these comparisons, scaled counts for all deregulated genes with log2FoldChange>1 and adjusted p-value <0.05 are shown in Figure 2 to provide an overview of effects and heterogeneity. Clusters were obtained with kmeans clustering using the superheat package. Gene lengths were obtained with EDAseq package used to provide gene length normalised read counts (RPKM) for supplementary files (Risso et al., 2011). Used ggplot for plotting except for heatmaps. Code for differential expression analysis available upon request.

Gene Set Enrichment Analysis (GSEA) was done using a GSEA software downloaded from the Broad Institute (Mootha et al., 2003; Subramanian et al., 2005). DESeq2 results were imported based on Wald Statistics as preranked lists and enrichment was calculated with the classic setting and 1000 permutations. Gene sets were downloaded from the Molecular Signature Database; mSigDB v7.2 (Chang et al., 2004; Hsiao et al., 2001; Liberzon et al., 2011). We performed GSEA against the entire C5: ontology gene sets (Table S3) and C8: cell type signature gene sets (Table S5) from mSigDB v7.2 as well as against selected gene sets representing cycling genes, liver-specific genes and genes enriched in developing human pancreas and liver. RNA-seq data generated in this study are available at the ArrayExpress database under the accession number E-MTAB-10715.

#### Public datasets

The ATAC-seq data (GSE149148, Geusz et al., 2021) and the ChIP-seq data (GSE61475, Tsankov et al., 2015) used in this study were downloaded from Gene Expression Omnibus (GEO) as processed data in bigwig and bed formats.

#### Statistical analysis and reproducibility

All statistics were performed using GraphPad Prism 8 software (GraphPad). Data sets with two groups having equal variances were analyzed by a two-tailed Student’s t test. For data with unequal variances, two-tailed Welch’s t tests were applied. Comparison of three or more groups was performed by one-way analysis of variance (ANOVA) followed by either Dunnett’s test using ctrl samples as reference, or Tukey’s test for comparison of all means. Statistics for differential expression of RNA-seq are provided as padj values from the DESeq2 analysis. *P* values are displayed in the figures and sample sizes are provided in the figure legends. Statistical significance is defined as p < 0.05 for qRT-PCR and GSEA data as well as FACS and IF image quantifications, while the DESeq2 analysis used an adjusted p-value (padj) cut-off set to 0.1 (default).

## References

Abu-Abed, S., Dolle, P., Metzger, D., Beckett, B., Chambon, P., and Petkovich, M. (2001). The retinoic acid-metabolizing enzyme, CYP26A1, is essential for normal hindbrain patterning, vertebral identity, and development of posterior structures. Genes Dev 15, 226–240.

Abu-Abed, S., Dolle, P., Metzger, D., Wood, C., MacLean, G., Chambon, P., and Petkovich, M. (2003). Developing with lethal RA levels: genetic ablation of Rarg can restore the viability of mice lacking Cyp26a1. Development 130, 1449–1459.

Ameri, J., Borup, R., Prawiro, C., Ramond, C., Schachter, K.A., Scharfmann, R., and Semb, H. (2017). Efficient Generation of Glucose-Responsive Beta Cells from Isolated GP2(+) Human Pancreatic Progenitors. Cell Rep 19, 36–49.

Arnold, S.J., and Robertson, E.J. (2009). Making a commitment: cell lineage allocation and axis patterning in the early mouse embryo. Nat Rev Mol Cell Biol 10, 91–103.

Bayha, E., Jorgensen, M.C., Serup, P., and Grapin-Botton, A. (2009). Retinoic Acid Signaling Organizes Endodermal Organ Specification along the Entire Antero-Posterior Axis. Plos One 4.

Belo, J.A., Bachiller, D., Agius, E., Kemp, C., Borges, A.C., Marques, S., Piccolo, S., and De Robertis, E.M. (2000). Cerberus-like is a secreted BMP and nodal antagonist not essential for mouse development. Genesis 26, 265–270.

Bhushan, A., Itoh, N., Kato, S., Thiery, J.P., Czernichow, P., Bellusci, S., and Scharfmann, R. (2001). Fgf10 is essential for maintaining the proliferative capacity of epithelial progenitor cells during early pancreatic organogenesis. Development 128, 5109–5117.

Bunt, J., Hasselt, N.E., Zwijnenburg, D.A., Hamdi, M., Koster, J., Versteeg, R., and Kool, M. (2012). OTX2 directly activates cell cycle genes and inhibits differentiation in medulloblastoma cells. Int J Cancer 131, E21–32.

Candia, A.F., Watabe, T., Hawley, S.H., Onichtchouk, D., Zhang, Y., Derynck, R., Niehrs, C., and Cho, K.W. (1997). Cellular interpretation of multiple TGF-beta signals: intracellular antagonism between activin/BVg1 and BMP-2/4 signaling mediated by Smads. Development 124, 4467–4480.

Chang, H.Y., Sneddon, J.B., Alizadeh, A.A., Sood, R., West, R.B., Montgomery, K., Chi, J.T., van de Rijn, M., Botstein, D., and Brown, P.O. (2004). Gene expression signature of fibroblast serum response predicts human cancer progression: similarities between tumors and wounds. PLoS Biol 2, E7.

Chung, W.S., Shin, C.H., and Stainier, D.Y. (2008). Bmp2 signaling regulates the hepatic versus pancreatic fate decision. Dev Cell 15, 738–748.

Costello,I., Nowotschin, S., Sun, X., Mould, A.W., Hadjantonakis, A.K., Bikoff, E.K., and Robertson, E.J. (2015). Lhx1 functions together with Otx2, Foxa2, and Ldb1 to govern anterior mesendoderm, node, and midline development. Genes Dev 29, 2108–2122.

Dang, L.T., Miao, Y., Ha, A., Yuki, K., Park, K., Janda, C.Y., Jude, K.M., Mohan, K., Ha, N., Vallon, M., et al. (2019). Receptor subtype discrimination using extensive shape complementary designed interfaces. Nat Struct Mol Biol 26, 407–414.

Deimling, S.J., and Drysdale, T.A. (2009). Retinoic acid regulates anterior-posterior patterning within the lateral plate mesoderm of Xenopus. Mech Dev 126, 913–923.

Deutsch, G., Jung, J., Zheng, M., Lora, J., and Zaret, K.S. (2001). A bipotential precursor population for pancreas and liver within the embryonic endoderm. Development 128, 871–881.

Dobin, A., Davis, C.A., Schlesinger, F., Drenkow, J., Zaleski, C., Jha, S., Batut, P., Chaisson, M., and Gingeras, T.R. (2013). STAR: ultrafast universal RNA-seq aligner. Bioinformatics 29, 15–21.

Fossat, N., Ip, C.K., Jones, V.J., Studdert, J.B., Khoo, P.L., Lewis, S.L., Power, M., Tourle, K., Loebel, D.A., Kwan, K.M., et al. (2015). Context-specific function of the LIM homeobox 1 transcription factor in head formation of the mouse embryo. Development 142, 2069–2079.

Galvin, K.E., Travis, E.D., Yee, D., Magnuson, T., and Vivian, J.L. (2010). Nodal signaling regulates the bone morphogenic protein pluripotency pathway in mouse embryonic stem cells. J Biol Chem 285, 19747–19756.

Gestri, G., Carl, M., Appolloni, I., Wilson, S.W., Barsacchi, G., and Andreazzoli, M. (2005). Six3 functions in anterior neural plate specification by promoting cell proliferation and inhibiting Bmp4 expression. Development 132, 2401–2413.

Geusz, R.J., Wang, A., Chiou, J., Lancman, J.J., Wetton, N., Kefalopoulou, S., Wang, J., Qiu, Y., Yan, J., Aylward, A., et al. (2021). Pancreatic progenitor epigenome maps prioritize type 2 diabetes risk genes with roles in development. Elife 10.

Goss, A.M., Tian, Y., Tsukiyama, T., Cohen, E.D., Zhou, D., Lu, M.M., Yamaguchi, T.P., and Morrisey, E.E. (2009). Wnt2/2b and beta-Catenin Signaling Are Necessary and Sufficient to Specify Lung Progenitors in the Foregut. Developmental Cell 17, 290–298.

Grapin-Botton, A. (2005). Antero-posterior patterning of the vertebrate digestive tract: 40 years after Nicole Le Douarin’s PhD thesis. International Journal of Developmental Biology 49, 335–347.

Hsiao, L.L., Dangond, F., Yoshida, T., Hong, R., Jensen, R.V., Misra, J., Dillon, W., Lee, K.F., Clark, K.E., Haverty, P., et al. (2001). A compendium of gene expression in normal human tissues. Physiol Genomics 7, 97–104.

Ip, C.K., Fossat, N., Jones, V., Lamonerie, T., and Tam, P.P. (2014). Head formation: OTX2 regulates Dkk1 and Lhx1 activity in the anterior mesendoderm. Development 141, 3859–3867.

Jennings, R.E., Berry, A.A., Gerrard, D.T., Wearne, S.J., Strutt, J., Withey, S., Chhatriwala, M., Piper Hanley, K., Vallier, L., Bobola, N., et al. (2017). Laser Capture and Deep Sequencing Reveals the Transcriptomic Programmes Regulating the Onset of Pancreas and Liver Differentiation in Human Embryos. Stem Cell Reports 9, 1387–1394.

Jung, J., Zheng, M., Goldfarb, M., and Zaret, K.S. (1999). Initiation of mammalian liver development from endoderm by fibroblast growth factors. Science 284, 1998–2003.

Kraus, M.R.C., and Grapin-Botton, A. (2012). Patterning and shaping the endoderm in vivo and in culture. Current Opinion in Genetics & Development 22, 347–353.

Lagutin, O.V., Zhu, C.C., Kobayashi, D., Topczewski, J., Shimamura, K., Puelles, L., Russell, H.R., McKinnon, P.J., Solnica-Krezel, L., and Oliver, G. (2003). Six3 repression of Wnt signaling in the anterior neuroectoderm is essential for vertebrate forebrain development. Genes Dev 17, 368–379.

Li, Y., Rankin, S.A., Sinner, D., Kenny, A.P., Krieg, P.A., and Zorn, A.M. (2008). Sfrp5 coordinates foregut specification and morphogenesis by antagonizing both canonical and noncanonical Wnt11 signaling. Genes Dev 22, 3050–3063.

Liberzon, A., Subramanian, A., Pinchback, R., Thorvaldsdottir, H., Tamayo, P., and Mesirov, J.P. (2011). Molecular signatures database (MSigDB) 3.0. Bioinformatics 27, 1739–1740.

Loh, K.M., Ang, L.T., Zhang, J., Kumar, V., Ang, J., Auyeong, J.Q., Lee, K.L., Choo, S.H., Lim, C.Y., Nichane, M., et al. (2014). Efficient endoderm induction from human pluripotent stem cells by logically directing signals controlling lineage bifurcations. Cell Stem Cell 14, 237–252.

Love, M.I., Huber, W., and Anders, S. (2014). Moderated estimation of fold change and dispersion for RNA-seq data with DESeq2. Genome Biol 15, 550.

Mahaddalkar, P.U., Scheibner, K., Pfluger, S., Ansarullah, Sterr, M., Beckenbauer, J., Irmler, M., Beckers, J., Knobel, S., and Lickert, H. (2020). Generation of pancreatic beta cells from CD177(+) anterior definitive endoderm. Nat Biotechnol 38, 1061–1072.

McLin, V.A., Rankin, S.A., and Zorn, A.M. (2007). Repression of Wnt/beta-catenin signaling in the anterior endoderm is essential for liver and pancreas development. Development 134, 2207–2217.

McMahon, J.A., and McMahon, A.P. (1989). Nucleotide sequence, chromosomal localization and developmental expression of the mouse int-1-related gene. Development 107, 643–650.

Miao, Y., Ha, A., de Lau, W., Yuki, K., Santos, A.J.M., You, C., Geurts, M.H., Puschhof, J., Pleguezuelos-Manzano, C., Peng, W.C., et al. (2020). Next-Generation Surrogate Wnts Support Organoid Growth and Deconvolute Frizzled Pleiotropy In Vivo. Cell Stem Cell 27, 840–851 e846.

Micallef, S.J., Janes, M.E., Knezevic, K., Davis, R.P., Elefanty, A.G., and Stanley, E.G. (2005). Retinoic acid induces Pdx1-positive endoderm in differentiating mouse embryonic stem cells. Diabetes 54, 301–305.

Mikels, A.J., and Nusse, R. (2006). Purified Wnt5a protein activates or inhibits beta-catenin-TCF signaling depending on receptor context. PLoS Biol 4, e115.

Molotkov, A., Molotkova, N., and Duester, G. (2005). Retinoic acid generated by Raldh2 in mesoderm is required for mouse dorsal Endodermal pancreas development. Developmental Dynamics 232, 950–957.

Monkley, S.J., Delaney, S.J., Pennisi, D.J., Christiansen, J.H., and Wainwright, B.J. (1996). Targeted disruption of the Wnt2 gene results in placentation defects. Development 122, 3343–3353.

Mootha, V.K., Lindgren, C.M., Eriksson, K.F., Subramanian, A., Sihag, S., Lehar, J., Puigserver, P., Carlsson, E., Ridderstrale, M., Laurila, E., et al. (2003). PGC-1alpha-responsive genes involved in oxidative phosphorylation are coordinately downregulated in human diabetes. Nat Genet 34, 267–273.

Nostro, M.C., Sarangi, F., Ogawa, S., Holtzinger, A., Corneo, B., Li, X., Micallef, S.J., Park, I.H., Basford, C., Wheeler, M.B., et al. (2011). Stage-specific signaling through TGFbeta family members and WNT regulates patterning and pancreatic specification of human pluripotent stem cells. Development 138, 861–871.

Ober, E.A., Verkade, H., Field, H.A., and Stainier, D.Y. (2006). Mesodermal Wnt2b signalling positively regulates liver specification. Nature 442, 688–691.

Onishi, K., and Zou, Y.M. (2017). Sonic Hedgehog switches on Wnt/planar cell polarity signaling in commissural axon growth cones by reducing levels of Shisa2. Elife 6.

Ortmann, D., Brown, S., Czechanski, A., Aydin, S., Muraro, D., Huang, Y., Tomaz, R.A., Osnato, A., Canu, G., Wesley, B.T., et al. (2020). Naive Pluripotent Stem Cells Exhibit Phenotypic Variability that Is Driven by Genetic Variation. Cell Stem Cell 27, 470–481 e476.

Pagliuca, F.W., Millman, J.R., Gurtler, M., Segel, M., Van Dervort, A., Ryu, J.H., Peterson, Q.P., Greiner, D., and Melton, D.A. (2014). Generation of functional human pancreatic beta cells in vitro. Cell 159, 428–439.

Piccolo, S., Agius, E., Leyns, L., Bhattacharyya, S., Grunz, H., Bouwmeester, T., and De Robertis, E.M. (1999). The head inducer Cerberus is a multifunctional antagonist of Nodal, BMP and Wnt signals. Nature 397, 707–710.

Poulain, M., and Ober, E.A. (2011). Interplay between Wnt2 and Wnt2bb controls multiple steps of early foregut-derived organ development. Development 138, 3557–3568.

Rankin, S.A., Kormish, J., Kofron, M., Jegga, A., and Zorn, A.M. (2011). A gene regulatory network controlling hhex transcription in the anterior endoderm of the organizer. Dev Biol 351, 297–310.

Rankin, S.A., McCracken, K.W., Luedeke, D.M., Han, L., Wells, J.M., Shannon, J.M., and Zorn, A.M. (2018). Timing is everything: Reiterative Wnt, BMP and RA signaling regulate developmental competence during endoderm organogenesis. Dev Biol 434, 121–132.

Rezania, A., Bruin, J.E., Arora, P., Rubin, A., Batushansky, I., Asadi, A., O’Dwyer, S., Quiskamp, N., Mojibian, M., Albrecht, T., et al. (2014). Reversal of diabetes with insulin-producing cells derived in vitro from human pluripotent stem cells. Nat Biotechnol 32, 1121–1133.

Rezania, A., Riedel, M.J., Wideman, R.D., Karanu, F., Ao, Z., Warnock, G.L., and Kieffer, T.J. (2011). Production of functional glucagon-secreting alpha-cells from human embryonic stem cells. Diabetes 60, 239–247.

Ribes, V., Fraulob, V., Petkovich, M., and Dolle, P. (2007). The oxidizing enzyme CYP26a1 tightly regulates the availability of retinoic acid in the gastrulating mouse embryo to ensure proper head development and vasculogenesis. Dev Dyn 236, 644–653.

Risso, D., Schwartz, K., Sherlock, G., and Dudoit, S. (2011). GC-content normalization for RNA-Seq data. BMC Bioinformatics 12, 480.

Rodriguez-Seguel, E., Mah, N., Naumann, H., Pongrac, I.M., Cerda-Esteban, N., Fontaine, J.F., Wang, Y., Chen, W., Andrade-Navarro, M.A., and Spagnoli, F.M. (2013). Mutually exclusive signaling signatures define the hepatic and pancreatic progenitor cell lineage divergence. Genes Dev 27, 1932–1946.

Rossi, J.M., Dunn, N.R., Hogan, B.L.M., and Zaret, K.S. (2001). Distinct mesodermal signals, including BMPs from the septum transversum mesenchyme, are required in combination for hepatogenesis from the endoderm. Genes & Development 15, 1998–2009.

Sanvitale, C.E., Kerr, G., Chaikuad, A., Ramel, M.C., Mohedas, A.H., Reichert, S., Wang, Y., Triffitt, J.T., Cuny, G.D., Yu, P.B., et al. (2013). A new class of small molecule inhibitor of BMP signaling. PLoS One 8, e62721.

Seiliez, I., Thisse, B., and Thisse, C. (2006). FoxA3 and goosecoid promote anterior neural fate through inhibition of Wnt8a activity before the onset of gastrulation. Dev Biol 290, 152–163.

Serls, A.E., Doherty, S., Parvatiyar, P., Wells, J.M., and Deutsch, G.H. (2005). Different thresholds of fibroblast growth factors pattern the ventral foregut into liver and lung. Development 132, 35–47.

Shim, W.J., Sinniah, E., Xu, J., Vitrinel, B., Alexanian, M., Andreoletti, G., Shen, S., Sun, Y., Balderson, B., Boix, C., et al. (2020). Conserved Epigenetic Regulatory Logic Infers Genes Governing Cell Identity. Cell Syst 11, 625–639 e613.

Sibbritt, T., Ip, C.K., Khoo, P.L., Wilkie, E., Jones, V., Sun, J.Q.J., Shen, J.X., Peng, G., Han, J.J., Jing, N., et al. (2018). A gene regulatory network anchored by LIM homeobox 1 for embryonic head development. Genesis 56, e23246.

Spence, J.R., Lauf, R., and Shroyer, N.F. (2011). Vertebrate intestinal endoderm development. Dev Dyn 240, 501–520.

Stafford, D., and Prince, V.E. (2002). Retinoic acid signaling is required for a critical early step in zebrafish pancreatic development. Current Biology 12, 1215–1220.

Stevens, M.L., Chaturvedi, P., Rankin, S.A., Macdonald, M., Jagannathan, S., Yukawa, M., Barski, A., and Zorn, A.M. (2017). Genomic integration of Wnt/beta-catenin and BMP/Smad1 signaling coordinates foregut and hindgut transcriptional programs. Development 144, 1283–1295.

Subramanian, A., Tamayo, P., Mootha, V.K., Mukherjee, S., Ebert, B.L., Gillette, M.A., Paulovich, A., Pomeroy, S.L., Golub, T.R., Lander, E.S., et al. (2005). Gene set enrichment analysis: a knowledge-based approach for interpreting genome-wide expression profiles. Proc Natl Acad Sci U S A 102, 15545–15550.

Tam, P.P., and Loebel, D.A. (2007). Gene function in mouse embryogenesis: get set for gastrulation. Nat Rev Genet 8, 368–381.

Tao, Q., Yokota, C., Puck, H., Kofron, M., Birsoy, B., Yan, D., Asashima, M., Wylie, C.C., Lin, X., and Heasman, J. (2005). Maternal wnt11 activates the canonical wnt signaling pathway required for axis formation in Xenopus embryos. Cell 120, 857–871.

Tsankov, A.M., Gu, H., Akopian, V., Ziller, M.J., Donaghey, J., Amit, I., Gnirke, A., and Meissner, A. (2015). Transcription factor binding dynamics during human ES cell differentiation. Nature 518, 344–349.

Wandzioch, E., and Zaret, K.S. (2009). Dynamic signaling network for the specification of embryonic pancreas and liver progenitors. Science 324, 1707–1710.

Yamamoto, A., Nagano, T., Takehara, S., Hibi, M., and Aizawa, S. (2005). Shisa promotes head formation through the inhibition of receptor protein maturation for the caudalizing factors, Wnt and FGF. Cell 120, 223–235.

Yao, J., and Kessler, D.S. (2001). Goosecoid promotes head organizer activity by direct repression of Xwnt8 in Spemann’s organizer. Development 128, 2975–2987.

Yasuoka, Y., Suzuki, Y., Takahashi, S., Someya, H., Sudou, N., Haramoto, Y., Cho, K.W., Asashima, M., Sugano, S., and Taira, M. (2014). Occupancy of tissue-specific cis-regulatory modules by Otx2 and TLE/Groucho for embryonic head specification. Nat Commun 5, 4322.

Zakin, L.D., Mazan, S., Maury, M., Martin, N., Guenet, J.L., and Brulet, P. (1998). Structure and expression of Wnt13, a novel mouse Wnt2 related gene. Mech Dev 73, 107–116.

Zaret, K.S. (2008). Genetic programming of liver and pancreas progenitors: lessons for stem-cell differentiation. Nat Rev Genet 9, 329–340.

