## Supplemental Methods and Figures for "TGF-β modulates cell fate in human ES cell-derived foregut endoderm by inhibiting multiple endogenous signaling pathways"

**Contents:**
**Key Resource Table**
**Supplemental Figures**

### KEY RESOURCES TABLE

| REAGENT or RESOURCE | SOURCE | IDENTIFIER |
|---|---|---|
| **Antibodies** | | |
| Goat polyclonal anti-PDX1 (WB 1:500) | R&D Systems | Cat# AF2419; RRID: AB_355257 |
| Mouse anti-NKX6.1 (IF 1:500) | DSHB | Cat# F55A12; RRID:AB_532379 |
| Goat polyclonal anti-OTX2 (IF, WB 1:500) | RnD Systems | Cat# AF1979; RRID: AB_2157172 |
| Rabbit anti-AFP (IF 1:500) | Agilent Dako | Cat# A0008; RRID: AB_2650473 |
| Mouse anti-AFP (WB 1:1000) | RnD | Cat# MAB1368; RRID: AB_357658 |
| Mouse anti-β-catenin (IF: 1:500) | BD Biosciences | Cat# 610154; RRID: AB_397555 |
| Goat anti human Fc IgG (IF, FACS: 1:100) | Jackson ImmunoResearch | Cat# 109-116-098; RRIS: AB_2337678 |
| Mouse monoclonal α-Vinculin (WB 1:5000) | Sigma-Aldrich | Cat# V9131; RRID: AB_477629 |
| DAPI (IF, FACS 1:2000) | Thermo Fisher Scientific | Cat# D1306; RRID: AB_2629482 |
| **Chemicals, peptides, and recombinant proteins** | | |
| DEF-CS | Takara Biosciences | Cat# Y30010 |
| MCDB131 | GIBCO/Thermo Fisher Scientific | Cat# 10372019 |
| BSA (fatty acid free) | Proliant Biologicals | Cat# 68700 |
| Sodium bicarbonate | Sigma-Aldrich | Cat# S6297 |
| D-(+)-Glucose solution | Sigma-Aldrich | Cat# G8769 |
| GlutaMAX | GIBCO/Thermo Fisher Scientific | Cat# 35050038 |
| L-Ascorbic Acid | Sigma-Aldrich | Cat# A4403 |
| Retinoic acid | Sigma-Aldrich | Cat# R2625 |
| SANT-1 | Selleckchem | Cat# S7092 |
| TPB (PKC activator) | Sigma-Aldrich | Cat# 565740 |
| T3 (3,3’,5-Triiodo-L-thyronine sodium salt) | Sigma-Aldrich | Cat# T6397 |
| ALK5 inhibitor II | Millipore | Cat# 616452 |
| LDN-193189 | Axon Medchem | Cat# 1509 |
| Zinc Sulfate | Sigma-Aldrich | Cat# Z0251 |
| Heparin | Sigma-Aldrich | Cat# H3149 |
| XAV-939 | Sigma-Aldrich | Cat# X3004 |
| SB431542 | Selleckchem | Cat# S1067 |
| SB-505124 | Sigma-Aldrich | Cat# S4696 |
| R115866 | Sigma-Aldrich | Cat# SML2092 |
| ITS-X | GIBCO/ Thermo Fisher Scientific | Cat# 51500056 |
| DAPT | Selleckchem | Cat# 2215 |
| PD98059 | Selleckchem | Cat# S1177 |
| CHIR-99021 | Axon Medchem | Cat# 2435 |
| IWP-L6 | Axon Medchem | Cat# 2212 |
| Recombinant Human TGF-β1 | Peprotech | Cat# 100-21 |
| Recombinant human HGF | Peprotech | Cat# 100-39 |
| Recombinant human EGF-L7 | Peprotech | Cat# 100-61 |

| | | |
|---|---|---|
| Recombinant human EGF | Peprotech | Cat# 100-15 |
| Recombinant human Activin A | Peprotech | Cat# 120-14E |
| Recombinant Human FGF7 (KGF) | Peprotech | Cat# 100-19 |
| Recombinant Human Wnt3a | R&D Systems | Cat# 5036-WN-010 |
| Recombinant Human Wnt11 | R&D Systems | Cat# 6179-WN-010 |
| Recombinant Human BMP4 | Peprotech | Cat# 120-05ET |
| Recombinant Human BMP6 | Peprotech | Cat# 120-06 |
| Recombinant Human RSPO3 | Peprotech | Cat# 120-44 |
| NGS-Wnt; DRPB-Fzd7/8 | Miao et al., 2020 | |
| TrypLE Express Enzyme (1x) | Thermo Fisher Scientific | Cat# 12604039 |
| Critical commercial assays | | |
| NEB NEXT Ultra II RNA Library Prep Kit | NEB | Cat# E7770 |
| mRNA magnetic isolation module for poly(A) purification | NEB | Cat# E7490 |
| NEB Next multiplex Oligos for Illumina | NEB | Cat# E7335 and E7500 |
| Click-IT EdU Alexa Fluor Flow Cytometry Assay Kit | Thermo Fisher Scientific | Cat# C10646 |
| Click-IT Plus EdU Cell Proliferation Kit for Imaging | Thermo Fisher Scientific | Cat# C10640 |
| Deposited data | | |
| RNA-Seq (raw and processed data) | This study | E-MTAB-10715 |
| Experimental models: cell lines | | |
| HUES4 PDX1$^{EGFP/+}$ | Ameri et al., 2017 | N/A |
| H1 (WA01) | WiCell | RRID: CVCL_9771 |
| Oligonucleotides | | |
| Primers: see Table S7 | | |
| Software and algorithms | | |
| Imaris version 9.2, 9.3, 9.5 | BitPlane | RRID: SCR_007370 |
| FCS Express version 6.04.0004 | De Novo Software | RRID: SCR_016431 |
| Bcl2Fastq Conversion Software | Illumina | RRID: SCR_015058 |
| R version 3.5.1 | https://r-project.org | N/A |
| R-studio version 1.1.463 | https://rstudio.com | N/A |
| DESeq2 | Love et al., 2014 | https://bioconductor.org/packages/release/bioc/html/DESeq2.html |
| STAR | Dobin et al., 2013 | https://github.com/alexdobin/STAR/archive/2.7.3a.tar.gz |
| EDASeq | Risso et al., 2011 | https://bioconductor.org/packages/release/bioc/html/EDASeq.html |
| GSEA software | Mootha et al., 2003; Subramanian et al., 2005 | https://www.gsea-msigdb.org/gsea/index.jsp |
| Microsoft Excel | Microsoft | RRID: SCR_016137 |
| Adobe Photoshop (CS6) | Adobe | RRID: SCR_014199 |
| Adobe Illustrator (CS6) | Adobe | RRID: SCR_010279 |
| GraphPad Prism version 8.2, 8.3, 9.1.2 | GraphPad | RRID: SCR_002798 |
| ZEN | Zeiss | RRID: SCR_013672 |
| Integrated Genome Viewer (IGV) v. 2.3.97 | Broad Institute | |
| Other | | |
| ATAC-seq data | Geusz et al., 2021 | GSE149148 |
| ChIP-seq data | Tsankov et al., 2015 | GSE61475 |

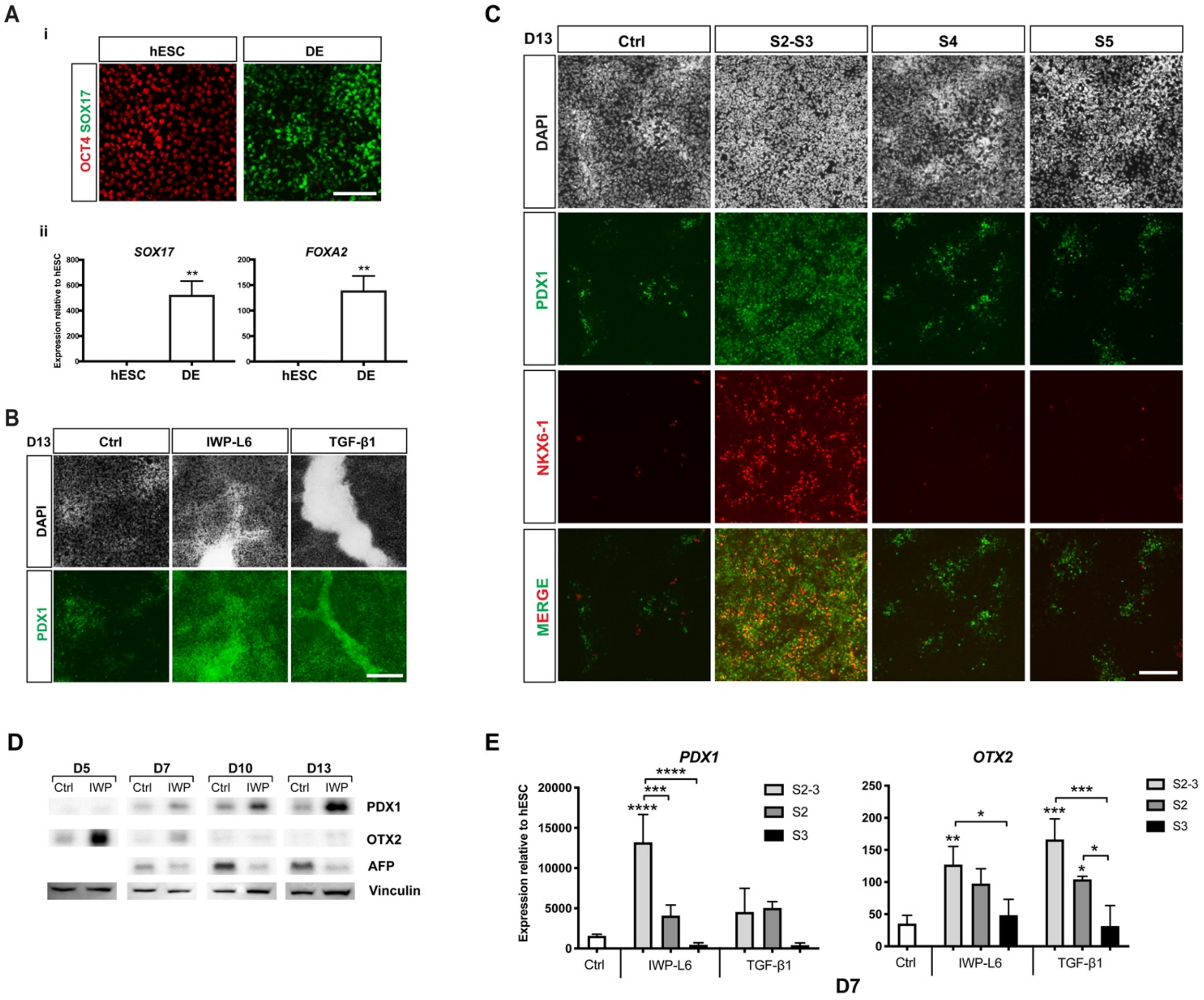


Figure S1

**Figure S1. Related to Figure 1. TGF-β1 and IWP-L6 anteriorize endoderm and promote pancreas over liver fate**

(A) IF staining for OCT4 (red) and SOX17 (green) protein (i) and expression of *SOX17* and *FOXA2* by qRT-PCR (ii) in *PDX1*$^{EGFP/+}$ HUES4 cells at the hESC and DE stages. Mean ± SD, N=3, ** $p < 0.005$. Scalebar, 50 μm.

(B) IF staining for PDX1 in D13 *PDX1*$^{EGFP/+}$ HUES4 cells treated with vehicle (Ctrl), IWP-L6 or TGF-β1 during S2-S5. Scalebar, 100 μm.

(C) IF staining for PDX1 and NKX6-1 in *PDX1*$^{EGFP/+}$ HUES4 cells at D13 treated with vehicle during S2-S5 (Ctrl) or IWP-L6 during S2-S3, S4 or S5. Scalebar, 50 μm.

(D) Western blot analysis of PDX1, OTX2 and AFP expression in *PDX1*$^{EGFP/+}$ HUES4 cells treated with vehicle (Ctrl) or IWP-L6 (IWP) during S2 (D5) or S2-S3 (D7, D10, D13). Vinculin is used as internal control.

(E) Expression of *PDX1* and *OTX2* by qRT-PCR in D7 *PDX1*$^{EGFP/+}$ HUES4 cells treated with vehicle (Ctrl), IWP-L6 or TGF-β1 during S2, S3 or S2-S3. Mean ± SD, N=3, * $p < 0.05$, ** $p < 0.005$, *** $p < 0.0005$, **** $p < 0.0001$.

A

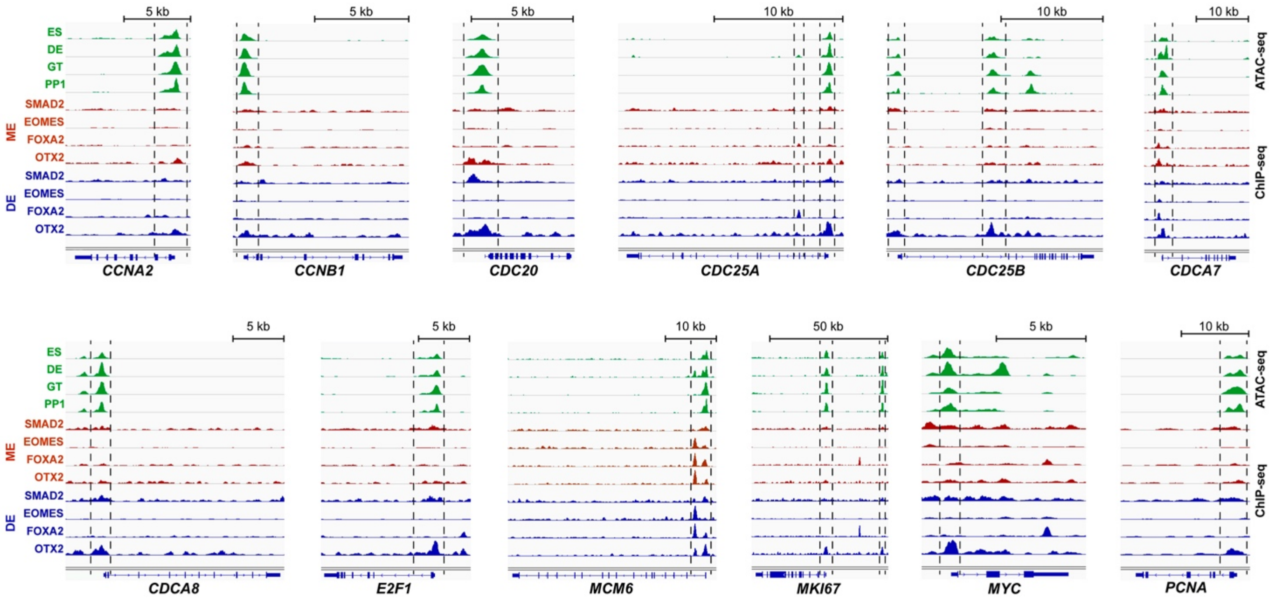


Figure S2

**Figure S2. Related to Figure 2. OTX2 occupies promoters of cell cycle related genes upregulated by TGF-β1 and IWP-L6**

(A) Signal tracks of ATAC-seq from Geusz et al. (2021) and SMAD2, EOMES, FOXA2 and OTX2 ChIP-seq data from Tsankov et al. (2015) at the loci of selected cell cycle related genes found upregulated by TGF-β1 and/or IWP-L6 in the RNA-seq analysis. Dashed lines are used to mark areas with chromatin accessibility at all stages and TF binding at the ME and/or DE stages. ES: ES cells; DE: definitive endoderm; GT: primitive gut tube; PP1: pancreatic progenitor 1, ME: mesendoderm.

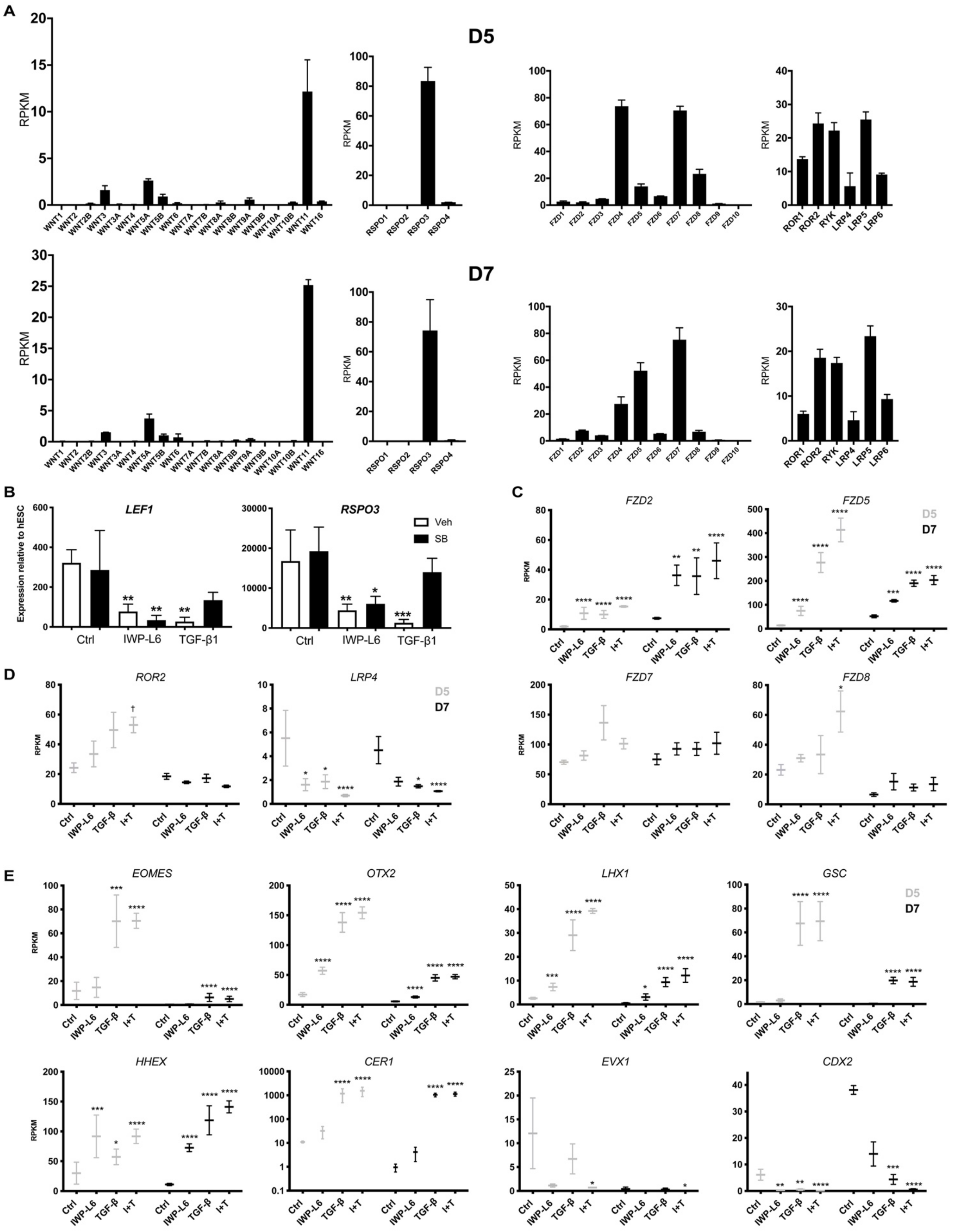

A
D5
D7
RPKM
WNT1
WNT2
WNT2B
WNT3
WNT3A
WNT4
WNT5A
WNT5B
WNT6
WNT7A
WNT7B
WNT8A
WNT8B
WNT9A
WNT9B
WNT10A
WNT10B
WNT11
WNT16
RSPO1
RSPO2
RSPO3
RSPO4
FZD1
FZD2
FZD3
FZD4
FZD5
FZD6
FZD7
FZD8
FZD9
FZD10
ROR1
ROR2
RYK
LRP4
LRP5
LRP6
B
Expression relative to hESC
LEF1
RSPO3
Veh
SB
Ctrl
IWP-L6
TGF-β1
C
FZD2
FZD5
D5
D7
D
ROR2
LRP4
FZD7
FZD8
E
EOMES
OTX2
LHX1
GSC
HHEX
CER1
EVX1
CDX2
Ctrl
IWP-L6
TGF-β
I+T


Figure S3

**Figure S3. Related to Figure 3. TGF-β1 activates an *OTX2*/*LHX1* GRN that antagonize Wnt signaling**

(A) Expression of WNT and RSPO ligand genes as well as FZD, ROR, RYK and LRP receptors and co-receptors by RNA-seq in D5 and D7 *PDX1*$^{EGFP/+}$ HUES4 control cells. Mean ± SD, N=3.

(B) Expression of Wnt target genes LEF1 and RSPO3 in D5 *PDX1*$^{EGFP/+}$ HUES4 cells in response to treatment with vehicle (Veh) or SB431542 (SB) in control (Ctrl), IWP-L6 or TGF-β1 treated cells. Mean ± SD, N=3, * $p < 0.05$, ** $p < 0.005$, *** $p < 0.0005$.

(C) Expression of *FZD* receptor genes by RNA-seq in D5 and D7 *PDX1*$^{EGFP/+}$ HUES4 cells treated with IWP-L6, TGF-β1 or IWP-L6 + TGF-β1 (I+T) during S2 (D5) and S2-3 (D7). Mean ± SEM, N=3, * padj < 0.05, ** padj < 0.005, *** padj < 0.0005, **** padj < 0.0001.

(D) Expression of *ROR2* and *LRP4* co-receptor genes by RNA-seq in D5 and D7 *PDX1*$^{EGFP/+}$ HUES4 cells treated with IWP-L6, TGF-β1 or IWP-L6 + TGF-β1 (I+T) during S2 (D5) and S2-3 (D7). † padj < 0.1, * padj < 0.05, **** padj < 0.0001.

(E) Expression of anterior markers from the *OTX2*/*LHX1*/*GSC* GRN and of posterior markers *EVX1* and *CDX2* by RNA-seq in D5 and D7 *PDX1*$^{EGFP/+}$ HUES4 cells treated with IWP-L6, TGF-β1 or IWP-L6 + TGF-β1 (I+T) during S2 (D5) and S2-S3 (D7). Mean ± SEM, N=3, * padj < 0.05, ** padj < 0.005, *** padj < 0.0005, **** padj < 0.0001.

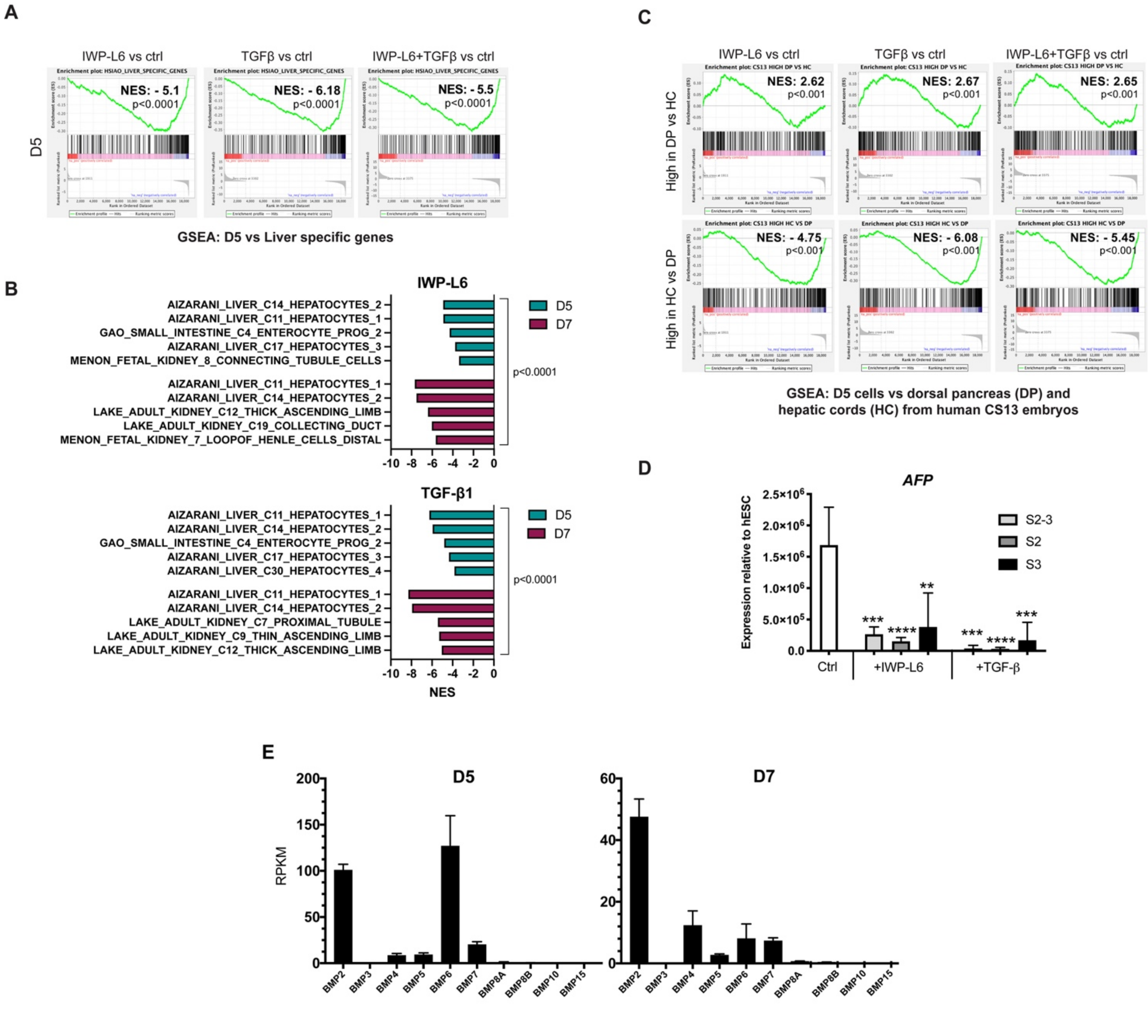


Figure S4

**Figure S4. Related to Figure 4. TGF-β1 promotes pancreas over liver fate by inhibiting BMP- and Wnt-signaling**

(A) GSEA plots comparing gene expression data from each treatment at D5 with a gene set for liver specific genes (Hsiao et al., 2001). Normalized Enrichment Score (NES) and p-values are shown.

(B) GSEA showing the top five negative NES scores when compared to RNA-seq data from D5 and D7 *PDX1*$^{EGFP/+}$ HUES4 cells treated with IWP-L6 or TGF-β1 to single-cell RNA-seq-based cell type-specific signatures.

(C) GSEA plots comparing gene expression data from each treatment at D5 with genes highly expressed in human Carnegie Stage (CS)13 dorsal pancreas (DP) vs hepatic cords (HC) and genes highly expressed in HC vs DP (Jennings et al., 2017).
(D) Expression of *AFP* by qRT-PCR in D7 *PDX1*$^{EGFP/+}$ HUES4 cells treated with vehicle (Ctrl), IWP-L6 or TGF-β1 during S2, S3 or S2-S3. Mean ± SD, N=3, ** $p < 0.005$, *** $p < 0.0005$, **** $p < 0.0001$.
(E) Expression of BMP ligand genes by RNA-seq in D5 and D7 *PDX1*$^{EGFP/+}$ HUES4 control cells. Mean ± SD, N=3.
