## Supplemental Tables for "TGF-β modulates cell fate in human ES cell-derived foregut endoderm by inhibiting multiple endogenous signaling pathways": Table_S7.docx

**STable S7. List of primers used in this study (related to STAR methods)**

| **Primers for qRT-PCR** |  |  |
| --- | --- | --- |
| Gene | Primer 1 (Fw) | Primer 2 (Rv) |
| *SOX17* | ACGCTTTCATGGTGTGGGCTAAG | GTCAGCGCCTTCCACGACTTG |
| *FOXA2* | GGAACACCACTACGCCTTCAAC | AGTGCATCACCTGTTCGTAGGC |
| *PDX1* | GAAGTCTACCAAAGCTCACGCG | GGAACTCCTTCTCCAGCTCTAG |
| *NKX6.1* | CCTATTCGTTGGGGATGACAGAG | TCTGTCTCCGAGTCCTGCTTCT |
| *OTX2* | GGAAGCACTGTTTGCCAAGACC | CTGTTGTTGGCGGCACTTAGCT |
| *CDX2* | ACAGTCGCTACATCACCATCCG | CCTCTCCTTTGCTCTGCGGTTC |
| *AFP* | CTTTGGGCTGCTCGCTATGA | TGGCTTGGAAAGTTCGGGTC |
| *SOX9* | AGGAAGCTCGCGGACCAGTAC | GGTGGTCCTTCTTGTGCTGCAC |
| *HNF6* | CAGGATGTGGAAGTGGCTTCAG | TGAACACCAGGCGGGACTTCTT |
| *HNF1β* | CCCAGCAAATCTTGTACCAGGC | ACCTCAGTGACCAAGTTGGAGC |
| *FABP1* | GGAGGAATGTGAGCTGGAGACA | TATGTCGCCGTTGAGTTCGGTC |
| *LEF1* | CTACCCATCCTCACTGTCAGTC | GGATGTTCCTGTTTGACCTGAGG |
| *RSPO3* | CCAGAAGGGTTGGAAGCCAACA | CCTTCTTCGTGCATGGACTCCA |
| *ID2* | TTGTCAGCCTGCATCACCAGAG | AGCCACACAGTGCTTTGCTGTC |
| *BMP6* | CCGACAACAGAGTCGTAATCGC | CTGCCATCCCAGGTCTTGGAAA |
| *WNT11* | CTGTGAAGGACTCGGAACTCGT | AGCTGTCGCTTCCGTTGGATGT |
| *RNF43* | GGTTACATCAGCATCGGACTTGC | ATGCTGGCGAATGAGGTGGAGT |
| *ROR2* | GTACGCATGGAACTGTGTGACG | AAAGGCAAGCGATGACCAGTGG |
| *ACTB* | CACCATTGGCAATGAGCGGTTC | AGGTCTTTGCGGATGTCCACGT |
